# The Balbiani body is formed by microtubule-controlled molecular condensation of Buc in early oogenesis

**DOI:** 10.1101/2022.03.11.484019

**Authors:** Swastik Kar, Rachael Deis, Adam Ahmad, Yoel Bogoch, Avichai Dominitz, Gal Shvaizer, Esther Sasson, Avishag Mytlis, Ayal Ben-Zvi, Yaniv M. Elkouby

## Abstract

Vertebrate oocyte polarity has been observed for two centuries and is essential for embryonic axis formation and germline specification, yet its underlying mechanisms remain unknown. In oocyte polarization, critical RNA-protein (RNP) granules delivered to the oocyte’s vegetal pole, are stored by the Balbiani body (Bb), a membraneless organelle conserved across species from insects to humans. However, the mechanisms of Bb formation are still unclear. Here, we elucidate mechanisms of Bb formation in zebrafish through developmental biomolecular condensation. Using super-resolution microscopy, live imaging, biochemical, and genetic analyses *in-vivo*, we demonstrate that Bb formation is driven by molecular condensation through phase-separation of the essential intrinsically disordered protein Bucky ball (Buc). Live imaging, molecular analyses, and FRAP experiments *in-vivo* reveal Buc-dependent changes in the Bb condensate’s dynamics and apparent material properties, transitioning from liquid-like condensates to a solid-like stable compartment. Furthermore, we identify a multi-step regulation by microtubules that controls Bb condensation: first through dynein-mediated trafficking of early condensing Buc granules, then by scaffolding condensed granules, likely through molecular crowding, and finally by caging the mature Bb to prevent overgrowth and maintain shape. These regulatory steps ensure the formation of a single intact Bb, which is considered essential for oocyte polarization and embryonic development. Our work offers insight into the long-standing question of the origins of embryonic polarity in non-mammalian vertebrates, support a paradigm of cellular control over molecular condensation by microtubules, and highlight biomolecular condensation as a key process in female reproduction.

## Introduction

Widely in animals and plants, gamete polarization lays the foundations for early embryonic development. In most vertebrates, including zebrafish, oocytes are polarized along an animal-vegetal axis which is established during oogenesis and is essential for embryonic development ^1-3^. RNA-protein (RNP) granules of embryonic patterning factors, including dorsal- and germline fate determinants localize to the oocyte and embryo vegetal pole, from which they later establish the global embryonic body axes and the germline lineage^1-3^. Those factors localize to the vegetal pole during oogenesis via a membraneless organelle, called the Balbiani body (Bb), which forms close to the oocyte nucleus and then translocates to the future vegetal pole^1-3^. Mutations that induce loss of the Bb or prevent its vegetal docking result in radially symmetrical eggs and early embryonic lethality^4-6^.

The Bb is conserved in oocytes from insects to humans^2,7-14^, where it forms in equivalent stages of oogenesis and exhibits similar dynamics^9,12,13,15^. However, despite its discovery dating back to 1845^9^, mechanisms of Bb formation are still unknown. In Zebrafish, during oocyte symmetry-breaking at the onset of meiosis, Bb granules first polarize around the centrosome by microtubules^12^. Bb granules then continue to aggregate around the centrosome within a nuclear indentation, called the nuclear cleft, and gradually assemble the mature compact Bb^12^ (cartoon in Figure 1A). However, the underlying mechanisms governing the gradual assembly of the Bb from a loose polarized aggerate to a mature compartment remain unknown.

**Figure 1.**
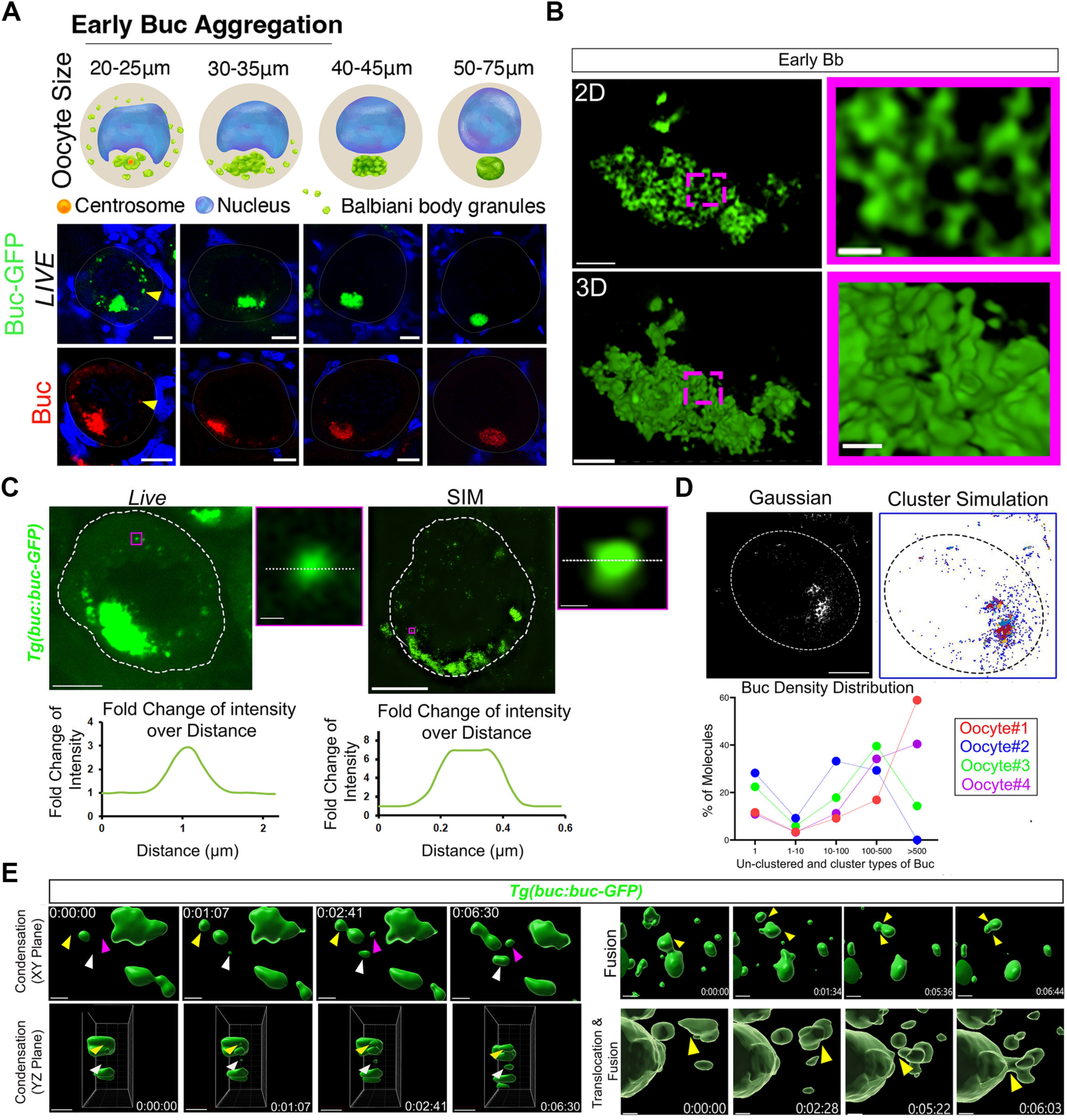
The Bb exhibit properties of a molecular condensate with phase-separation of Buc. **A.** Top: Bb formation during oocyte development, indicating oocyte stages and sizes. Bottom: Live imaging of transgenic Buc-GFP (green, top) and antibody labeling of endogenous Buc (red, bottom) detect small granules (arrowheads) that are not localized to the main aggregate, specifically during early Bb formation, and not in later formation or in the mature Bb stages (two right panels). **B.** Fine structure of the forming Bb aggregate at early cleft stages, as detected by super-resolution SIM imaging shows early network of Buc-GFP (green) which appears like loose viscoelastic material. 2D (top) and 3D (bottom) images are shown. Magenta-boxed zoom in images are magnifications of the dashed boxes in the left panels. Scale bars are 3 μm and 0.5 μm in the zoom out and zoomed in images, respectively. Images are snapshots from Videos S1. **C.** 3D analysis of individual Buc-GFP granules (green) by live confocal- (left) and fixed super-resolution SIM- (right) imaging shows granules with roundish irregular geometry. Zoom in images (right) are example individual granules magnified from boxed regions in oocyte images (outline; left). Scale bars are 3 μm and 0.5 μm in the zoom out and zoomed in images, respectively. Measurements of Buc-GFP intensities of granules and their immediately neighboring cytoplasm (dashed white lines) are plotted (bottom). Representative granules and plots are shown. **D.** dSTORM cluster analysis of Buc, showing Gaussian presentation (left) and cluster simulation (right) images of a representative oocyte (outline). Individual simulated clusters are color coded. Scale bar is 10 μm. The distribution of Buc molecules from four representative oocytes is plotted (bottom). Buc molecules are detected as un-clustered individual molecules, or in distinct clusters of different sizes (binned), and the percentage of molecules in each category is shown. Data from each oocyte are color coded and connected with a line. **E.** Live time-lapse imaging of Buc-GFP granules (green) in ovaries exhibit liquid droplet-like dynamics, including: left - granules that condense in real time (top is XY view, bottom is YZ view, color-coded arrowheads tracking individual granules), top right – granules that fuse with other granules (arrowheads), and bottom right - granules that translocate to and fuse with the main aggregates (arrowheads). Scale bars are 2 μm. Time (hh:mm:ss) is indicated. Images are snapshots from Videos S2. See also Videos S1-2.

Molecular condensates in cells are complexes that phase-transition to separate from the cytoplasm, forming membraneless compartments with specialized content and functions (reviewed in^16-20^). RNP granules form molecular condensates^21,22^, in somatic cells^18^, and specifically in germ cells and early embryos^17,23^. Foundational principles of RNP condensation were derived from *in-vitro* studies, cell cultures, and *in-vivo* research in *C. elegans* and *Drosophila* embryos^16-20,24-30^. Intrinsically disordered regions (IDRs) in proteins^31^, along with ‘sticker’ and ‘spacer’ regions in condensate components^32,33^, facilitate phase-separation through multivalent interactions. These interactions are generally self-assembling (e.g.,^34-37^), but can be influenced by temperature and pH^16-20,38,39^, RNA binding^22,30,40^, and post-translational modifications^32,33,41-43^. Condensates can shift between material states over time^44,45^, from liquid-like to solid-like^39^. In cells, they likely exhibit viscoelastic properties, forming through phase-separation combined with other processes (termed phase-separation++, PS++)^44,45^. Cell environments, including other molecules, cytoskeletal elements, and organelles, likely also shape their viscoelastic properties and dynamics.

Several observations suggest that the Bb may form through molecular condensation. First, the Bb contains RNP granules, and its essential protein, Bucky ball (Buc)^10,46^, is an intrinsically disordered protein (IDP)^46,47^. Second, our previous research showed that Buc is not required for symmetry breaking, but is essential for maintaining the integrity of the Bb aggregate in the cleft^12^, indicating a direct role in Bb condensation. Finally, we observed that the initial aggregate becomes progressively denser and more compact in forming the mature compartment (Figure 1A). Similarly, in *Xenopus*, the mature Bb was described as a prion-like aggregate with amyloid β-sheets^10,48^. Prion-like aggregates occupy the solid end of the condensation spectrum^10,39,49^, hinting at possible Buc-driven condensation dynamics during early Bb formation. Here, we investigate the mechanisms of Bb formation, from initial aggregation post-symmetry breaking to maturation.

## Results

### Early dynamic formation of the Balbiani body

We examined Buc dynamics in early oogenesis using antibody labeling of the endogenous Buc protein in *wt* ovaries, and by live imaging of transgenic Buc-GFP in *Tg(buc:Buc-GFP-buc3’utr)* ovaries^4^. *Tg(buc:Buc-GFP-buc3’utr)* consists of the *buc* genomic locus, recapitulating endogenous Buc-GFP protein localization and function^4^. Based on prior studies of Bb formation^12^, we focused on early, mid, and late cleft stages, as well as mature Bb stages (Figure 1A).

In late cleft and mature Bb stages we observed a single localized aggregate of either Buc or Buc-GFP (n=34 oocytes, 5 ovaries). By contrast, early and mid-cleft stages showed additional smaller granules of Buc and Buc-GFP outside the main aggregate (n=50 oocytes, 6 ovaries) (Figure 1A). These smaller granules could localize and contribute to the main aggregate, indicating dynamic Buc behavior during early Bb development. Three-dimensional super-resolution structured illumination microscopy (SIM) on whole Buc-GFP ovaries (Figure 1B) showed that early Buc-GFP aggregates appeared as a loose network that seemed like viscoelastic material (n=15 oocytes, 5 ovaries; Figure 1B, Video S1), typical of molecular condensates^19,44,45,50^. Given Buc’s potential dynamics and apparent material properties, we explored whether Bb forms through Buc-driven molecular condensation.

### Balbiani body formation is mediated by molecular condensation and Buc phase-separation

Phase-separation is defined as density transition, where molecules in a system (e.g., a cell’s cytoplasm) can exist in co-existing phases: a low density (dilute) phase and high density (condensed) phase^33,44,45^. Buc and Buc-GFP signals in oocytes exhibited low fluorescent intensity in the cytoplasm, along with high fluorescent intensity within granules (Figure 1A, C), suggesting that they represent phase-separated entities^44,45^. We analyzed individual granules in 3D through live confocal (n=50 granules, 12 oocytes, 5 ovaries), and SIM (n=35 granules, 8 oocytes, 5 ovaries) imaging. Detected granules were generally roundish but irregular (Figure 1C), similar to stress granules^51^. Condensates are often spherical due to liquid-like surface tension^19,50^, but stress granules in cells are typically irregular, non-spherical, and have viscoelastic properties with varying interfacial tensions and bending rigidities^44,45,51^. Fluorescent intensity measurements of Buc-GFP granules confirmed a 2.56+1.03 to 6.38+2.66-fold higher Buc-GFP intensity within granules, compared to neighboring cytoplasm (Figure 1C).

In cells or cell-free systems, density transition is often analyzed by measuring the refractive index as a proxy for material density^52,53^. Given the challenge of measuring refractive index in dense tissues, we directly measured Buc molecule density using direct stochastic optical reconstruction microscopy (dSTORM) super-resolution microscopy with single-molecule detection at 20 nm resolution (Figure 1D; Methods). Cluster analysis on Buc single molecules in whole oocytes (Methods), revealed isolated singles, small clusters (<10 molecules), or larger clusters with dozens to hundreds of molecules (n=25,546 signals from 4 ovaries; Figure 1D), indicating non-homogeneous Buc distribution.

To capture a higher temporal resolution of Buc dynamics, we conducted live time-lapse imaging of Buc-GFP in whole ovaries (Figure 1E; Video S2; Methods). We analyzed granule dynamics in 3D through Z-stacks of entire oocytes over time (Figure 1E; Video S2; Methods). Buc-GFP granules were observed outside the cleft and around the main aggregate, revealing three types of dynamics (n=25 oocytes, 5 ovaries): ***1)*** Granules appearing “de-novo” in the cytoplasm, with increasing intensity suggesting real-time condensation (Figure 1E, Video S2); ***2)*** Granules fusing with other granules or the main aggregate (Figure 1E, Video S2); and ***3)*** Granules translocating to and fusing with the main aggregate (Figure 1E, Video S2). Fusion ability is a defining characteristic of molecular condensates exhibiting liquid droplet-like behavior. While condensates in cells are typically not simple fluids^44,45,51^, the observed condensation and fusion of Buc-GFP granules resemble properties of liquid-like condensates.

We investigated if Buc granules form a non-soluble phase under physiological conditions. Stage-specific oocytes from Buc-GFP transgenic ovaries were isolated based on size^7,47,54^. In mature Bb-stage oocytes isolated this way, Buc-GFP displayed a clear signal in the mature Bb (Figure 2A). We then lysed oocytes from all Bb formation stages, and separated soluble supernatant and insoluble pellet fractions by centrifugation (Figure 2A). Western blot analysis showed that Buc-GFP was present specifically in the non-soluble pellet fraction and absent in the soluble supernatant (Figure 2B). Adding 0.5-1.5M L-Arginine (a solubilizing agent^55^) to the lysis buffer shifted Buc-GFP to the soluble supernatant in a dose-dependent manner (Figure 2B), confirming that Buc-GFP’s presence in the pellet fraction results from its insolubility. Thus, under physiological conditions, Buc-GFP exists as part of insoluble complexes.

**Figure 2.**
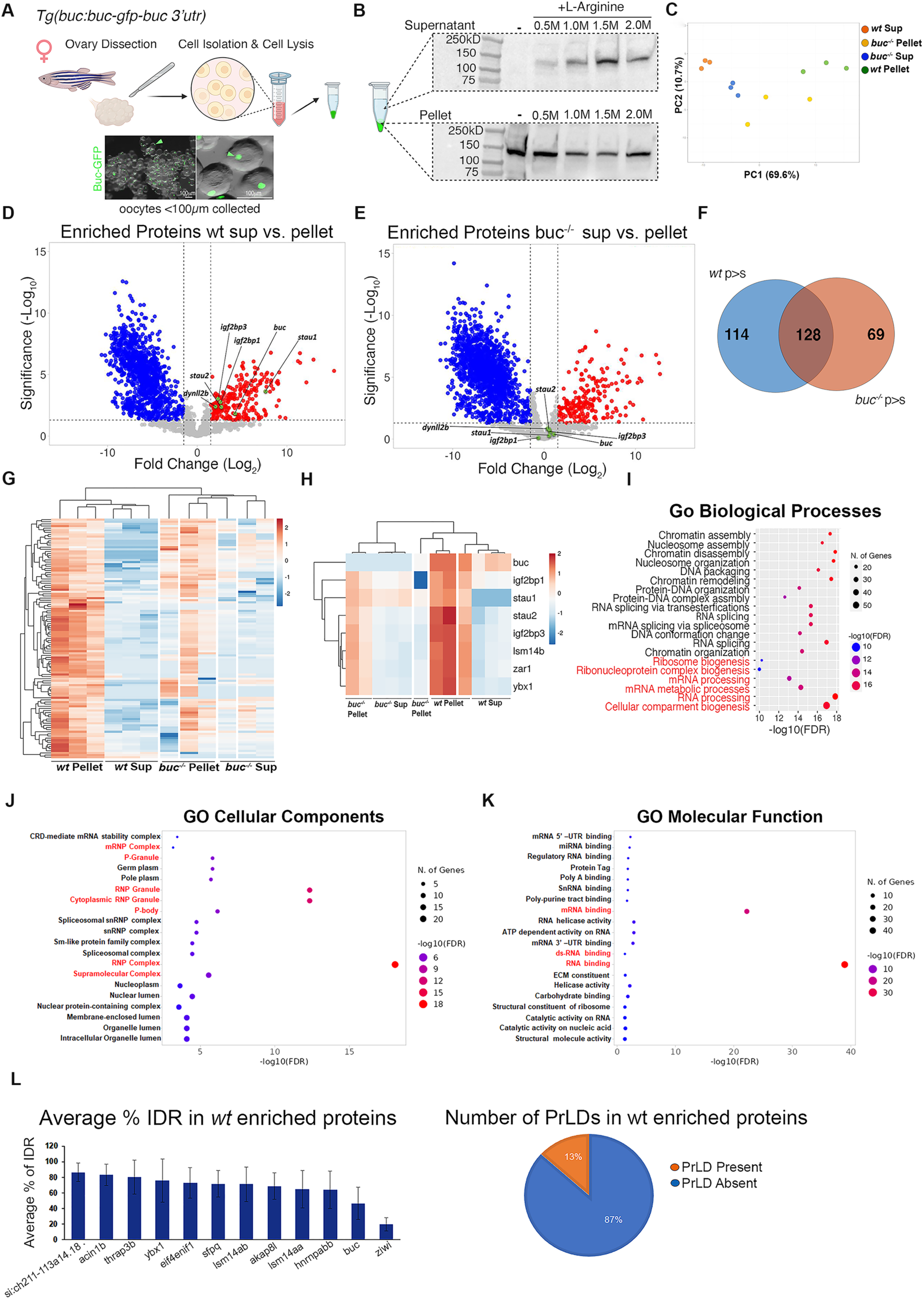
Bb RNAs form insoluble complexes in a Buc dependent manner. **A.** Experimental design: Ovaries are dissected from Buc-GFP fish, oocytes at all Bb stages (all sizes < 100 μm in diameter) are isolated, and lysed. Oocyte lysate is centrifuged, separating a soluble supernatant fraction and a non-soluble pellet fraction. **B.** Western blot analysis shows Buc-GFP detection in the pellet, but not supernatant fraction. Addition of L-Arg to the lysis buffer shifts Buc-GFP to the supernatant fractions in a dose-dependent manner until saturation at 1.5 M. **C-H.** Mass-spectrometry analyses of pellet and supernatant fractions from wt and *buc^-/-^*ovaries. PCA analysis (C) Volcano plots (D-E) Venn diagram (F) showing enrichment of proteins specifically in the pellet over the supernatant fraction in the wt samples (D), but not in *buc^-/-^* samples (E). **G-H.** Heatmaps with hierarchical clustering of the proteins that are specifically enriched in the pellet over supernatant samples in wt, but loose enrichment in *buc^-/-^* samples (F, see also C-D), including select known Bb proteins (G). **I-K.** GO analyses of Buc-dependent pellet enriched proteins identifies multiple RNA biogenesis and processing processes (red highlight). **L.** Left: Top ten Buc-dependent insoluble Bb proteins that contain substantial (∼65-85%) IDR sequences. Buc (∼45%) and Ziwi (∼20%) are shown as positive and negative IDR control proteins, respectively. Right: Percentage of Buc-dependent insoluble Bb proteins with or without a predicted Prion-like domain (PrLD). See also Table S1.

To determine if insoluble Buc-GFP complexes represent Bb complexes, we analyzed pellet and supernatant fractions from oocytes isolated from *wt* and *buc*^-/-^ ovaries using mass spectrometry. We expected Bb proteins in the insoluble pellet to be Buc-dependent, with Bb-insoluble complexes becoming more dissolved in *buc*^-/-^ samples due to Bb loss. We hypothesized that Bb resident proteins would be enriched in the *wt* pellet but not in the *buc*^-/-^ pellet. Principal component analysis (PCA) showed clustering of *wt* pellet triplicates, separate from wt supernatant triplicates (Figure 2C). As anticipated, *buc*^-/-^ supernatant samples clustered near *wt* supernatant clusters (Figure 2C). However, *buc*^-/-^ pellet samples did not cluster with *wt* pellets and were more dispersed, indicating that a significant cohort of *wt* pellet-enriched proteins lost their enrichment in *buc*^-/-^ pellet samples (Figure 2C).

We identified 114 proteins specifically enriched in *wt* pellet samples but not in *buc*⁻ pellet samples (Figure 2D-G; Table S1). These *wt* pellet-enriched proteins included known Bb-associated proteins, such as Buc, Zarl, Ybx, Staufen1/2, and Igf2bp3^47^ (Figure 2D-H), validating the presence of Bb complexes in the insoluble *wt* pellet fraction. In contrast, these proteins were detected at similar levels in both *buc*^-/-^ supernatant and pellet samples, indicating a shift to the soluble fraction following Buc and Bb loss (Figure 2D-H). Thus, the insolubility of Bb proteins depends on Buc and/or their Bb association.

Gene ontology analysis revealed that many Buc-dependent insoluble Bb complexes are associated with RNA biogenesis and processing, and RNP granules (Figure 2I-K). Most enriched proteins were predicted to contain IDRs (Figure 2L), with 13% containing prion-like domains (PLD) similar to Buc and *Xenopus* XVelo (Figure 2L). These results biochemically confirm the insoluble nature of Buc-dependent Bb RNP complexes *in-vivo*. In summary, our data support that Bb formation is mediated by molecular condensation driven by Buc phase separation and/or PS++ (hereafter referred to inclusively as phase separation for simplicity).

### Developmental changes in Buc dynamics and apparent material properties during Bb formation

The mature Bb appears more compact (Figure 3E), and in *Xenopus* has been described as an inert solid structure^10,48^. The viscoelastic properties of condensates change over time^44,45^, and transitioning toward a solid-like state is thought to be due to increased multivalent interactions^19,44,45,50^. We investigated possible developmental changes in Buc dynamics and in the apparent material properties of the Bb.

**Figure 3.**
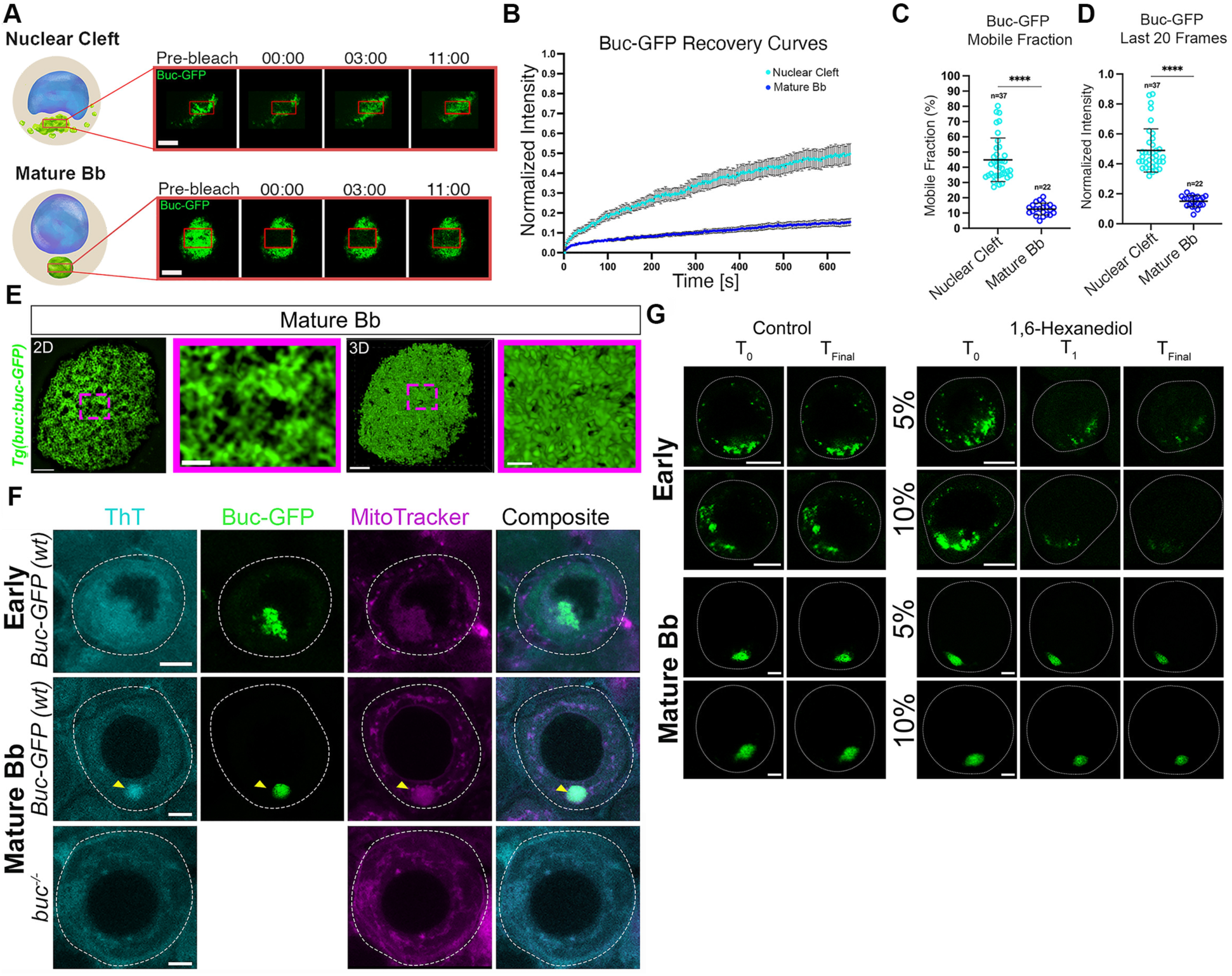
Developmental changes of Buc dynamics and apparent viscoelastic properties of the Bb. **A-D.** FRAP analyses of Buc-GFP turnover in in whole ovaries, in the early forming Bb in the nuclear cleft (top) and in the mature Bb (bottom) stages. Images in A are snapshots at indicated timepoints (minutes) from representative Video S3. Recovery plots (B), as well as quantification by mobile and immobile (C) fraction calculations, and by intensity at last 20 frames (D) are shown. All bars are Mean ± SD. See also Figure S1A-B. **E.** Fine structure of the mature Bb, as detected by super-resolution SIM imaging shows a dense and rugged network of Buc-GFP (green) compared to early stages in Figure 1B. 2D (left) and 3D (right) images are shown. Magenta-boxed zoom in images are magnifications of the dashed boxes in the left panels. Scale bars are 3 μm and 0.5 μm in the zoom out and zoomed in images, respectively. Images are snapshots from Videos S5. **F.** Live detection in Buc-GFP ovaries (phenotypically wt) of presumptive amyloid β-sheets (ThT, cyan) in mature Bb (arrowheads, middle), as labeled by transgenic Buc-GFP (green) and mitochondria (MitoTracker, magenta). ThT is not detected above background levels in the early Buc-GFP-positive, mitochondria-enriched forming Bb (top). At mature Bb stages in *buc^-/-^*ovaries, the Bb fails to form as indicated by MitoTracker, and ThT is not detected above background levels (bottom). Transgenic Buc-GFP protein is not expressed in the *buc* mutant line. Scale bars are 10 μm. See also Figure S2A. **G.** Treatment of live cultured Buc-GFP (green) ovaries with vehicle control (left) or with 5% or 10% 1,6-Hexanediol (right panels). *T_0_* shows oocytes prior to treatment. Each control oocyte (*T_0_-T_Final_*) corresponds to the oocytes in the 5% or 10% Hexanediol treated ovaries, respectively. Images of early (top), and mature Bb (bottom) are shown. Scale bars are 10 μm. See also Videos S3, 5, Figures S1-2.

To assess Buc developmental dynamics, we measured its turnover during Bb formation using fluorescence recovery after photobleaching (FRAP) assays in live whole-mount Buc-GFP ovaries. We recorded Buc-GFP recovery in early-mid cleft and mature Bb stage oocytes (Figures 3A-D, S1A, Video S3; Methods). MitoTracker vital dye was used as a cell viability proxy, ruling out artifacts from cell damage in FRAP experiments (Figure S1B). All recovery curves are shown in Supp. Figure 1A, with pooled data for cleft and mature Bb stage oocytes plotted in Figure 3B (n=37 cleft stage oocytes, 8 ovaries; n=22 mature Bb stage oocytes, 6 ovaries).

In early-mid cleft stages, Buc-GFP displayed significant recovery (∼40-50%) within the recovery time, as early as 3 minutes post-photobleaching (Figures 3A-B, S1A, Video S3), supporting the dynamic nature of the early Bb condensate. By contrast, mature Bb showed minimal recovery after photobleaching, with almost no Buc-GFP signal in the bleached area (Figure 3A-B, Video S3). Recovery curves indicated only ∼10% recovery over ∼600 seconds (n=22 mature Bb stage oocytes, 6 ovaries; Figure 3A-B, S1A, Video S3). Calculated Buc-GFP mobile fractions^56-58^ (Methods) were 44.88±14.29% in cleft stage oocytes and only 12.57±3.86% in mature Bb stages (Figure 3C). Consistently, relative Buc-GFP intensity during the final 20 frames post-bleaching (Methods) was 0.49±0.14 at cleft stages, dropping to 0.15±0.03 in mature Bb stages (Figure 3D).

As controls, we performed FRAP experiments with two additional transgenically expressed proteins: free GFP [*Tg(vasa:GFP)*] and H2A-GFP [*Tg(h2a:H2A-GFP)*]. Bleaching cytoplasmic free GFP resulted in fast and high recovery, as expected^59^ (Figure S1C-I, Video S4). Nuclear H2A-GFP showed low and slow recovery, consistent with previous findings^60^ (Figure S1J-P, Video S4). Both GFP and H2A-GFP exhibited similar recovery dynamics in early cleft (n=11 oocytes, 5 ovaries for each) and mature Bb stage oocytes (n=11 oocytes, 5 ovaries for each; Figure S1C-P, Video S4). These controls indicate that the distinct recovery rates observed for Buc-GFP are specific to Buc’s biological properties, confirming reduced turnover and increased stability in the mature Bb.

While FRAP alone cannot determine material properties^44,50,61^, the reduced dynamics in the mature Bb can point to potential development of more solid-like viscoelastic properties. We imaged mature Bbs in oocytes and ovaries from our SIM experiments in Figure 3E. In contrast to the loose network of Buc-GFP in early-mid cleft stages (Figure 1B, Video S1), in the mature Bb, this network appeared more dense and rugged (n= 11 oocytes, 5 ovaries; Figure 3E, Video S5), similar to the *Xenopus* mature Bb. The *Xenopus* Bb contains prion-like amyloid β-sheets^10,48^, which form highly ordered fibrillar aggregates in a solid-like state^62,63^.

To detect presumptive amyloid β-sheets in zebrafish ovaries, we used Thioflavin T (ThT), a dye also used for *Xenopus* Bb^49^. We compared early cleft-stage and mature Bb condensates. In cleft stage oocytes, ThT showed only background levels, unrelated to the forming Bb marked by Buc-GFP and Bb mitochondria (n=46 oocytes, 4 ovaries; Figure 3F). In mature Bb stage oocytes, however, ThT was specifically detected in Buc-GFP-positive, mitochondria-enriched Bb (n=62 oocytes, 4 ovaries; Figure 3F), confirmed by co-labeling with endogenous Buc and the Bb marker DiOC6^7,12^ (n=73 oocytes, 5 ovaries; Figure S2A). ThT signals were absent in *buc^-/-^* ovaries (n=51 oocyte, 4 ovaries), where Bb loss was verified with Mitotracker (Figure 3F), and co-labeling with Buc and DiOC6 (Figure S2A). The specific Buc-dependent detection of ThT signals in the mature Bb suggests that amyloid sheets form during later Bb development.

We tested Bb resistance to 1,6-Hexanediol, which disrupts early, reversible IDR interactions but not solid-like condensates^64^. Treating live ovaries with 5% or 10% Hexanediol disrupted Buc condensates at early cleft stages within 30 minutes, nearly abolishing them by 90 minutes (n=63 oocytes, 8 ovaries; Figure 3G). Fixed-ovary validation showed both endogenous Buc and *dazl* mRNA in Bb granules were sensitive to Hexanediol at both concentrations (n=4 control ovaries, n=5 treated ovaries per concentration; Figure S2B), confirming Hexanediol dissolves early Bb condensates. In contrast, Buc condensates in mature Bbs were highly resistant to both 5% and 10% Hexanediol, even after 90 minutes of treatment (n=56 oocytes, 7 ovaries; Figure 3G). This suggests that IDR interactions in the mature Bb are irreversible within the observed timeframe. The mature Bb’s insensitivity to Hexanediol may stem from limited penetration into dense complexes. The detection of Buc-dependent β-sheets and reduced Buc turnover in FRAP, however, support solid-like properties, likely due to saturated interactions among Buc and Bb constituents. Non-specific effects of 10% Hexanediol have been noted in cell cultures and yeast^65^, but consistent results with 5% treatment and likely reduced effective concentration in whole ovaries compared to cell culture and yeast, indicate minimal artifact in our setup.

In summary, our data reveal Bb formation dynamics. It is possible that early Buc granules already contain small solid-like clusters and then aggregate into the main condensate. However, based on our data indicating liquid-like dynamics of early granules from live imaging, Buc-dependent late presumptive amyloid formation, and differential sensitivity to Hexanediol, we favor the following model: Buc initially phase-transitions Bb RNP granules into small, liquid-like condensates that fuse into the main cleft condensate. In the cleft, Buc-dependent modifications (likely by saturating interactions with Bb proteins/RNA) lead to amyloid β-sheets and the solid-like properties of the mature Bb. A combined model is also possible, whereby “solidifying” Buc-dependent modifications begin in granules as they join the main condensate where they continue progressively.

### Microtubules are required for Buc turnover in the early forming Bb condensate

Control over molecular condensation is critical for function. In *buc*^-/-^ oocytes, Bb mRNAs are randomly distributed, leading to non-polarized eggs and embryonic lethality^5^. This led us to investigate what regulates Buc localization and turnover in condensates. We previously showed microtubules are essential for Buc localization during symmetry breaking^12^, and detected microtubules around the centrosome in the nuclear cleft^12^. To further characterize this, we used transgenic *Tg(βAct:EMTB-3GFP)* ovaries or α-tubulin labeling (Figures 4A-C, S3).

**Figure 4.**
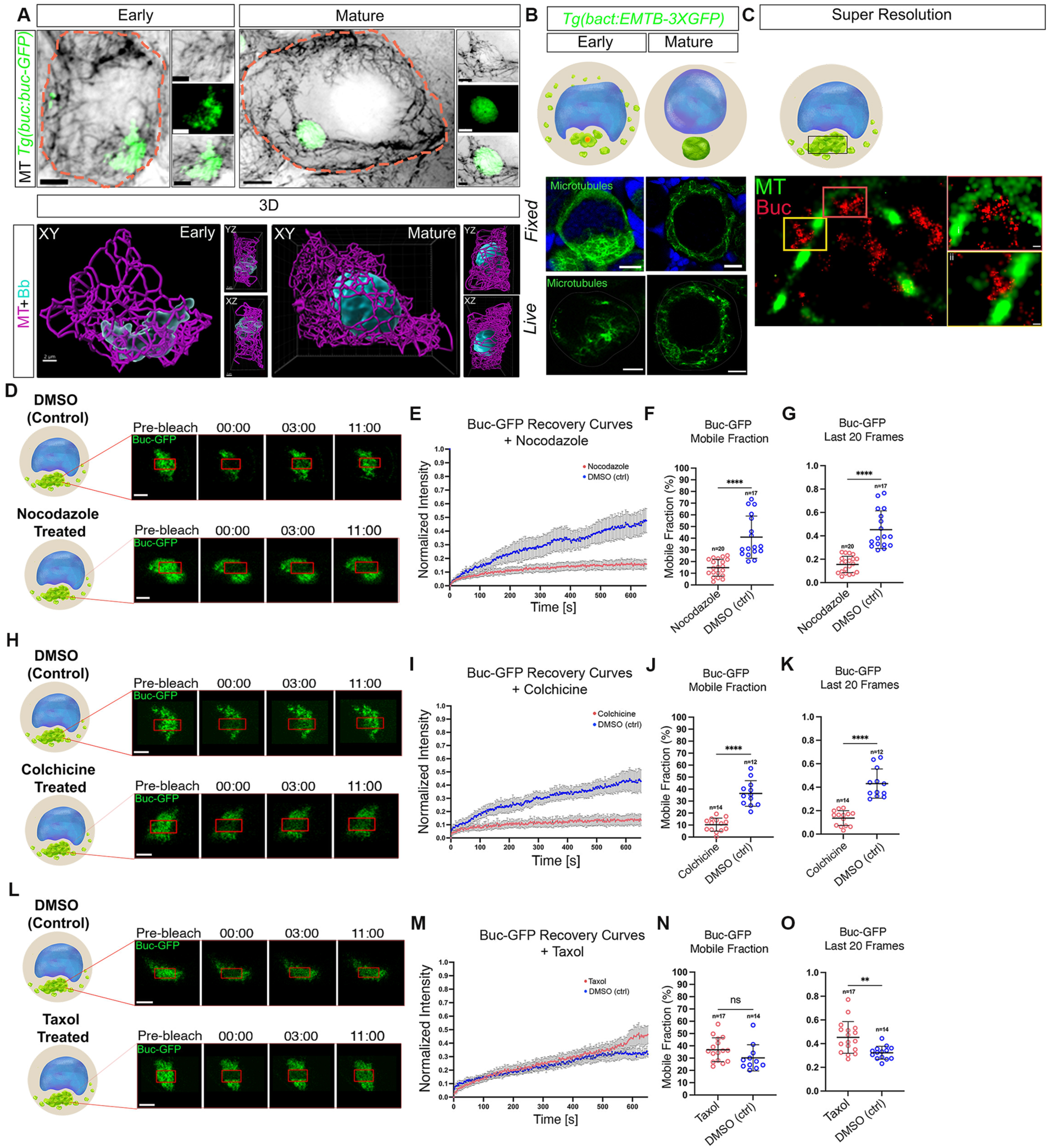
Buc turnover in the forming Bb condensate requires dynamic organization of microtubules. **A.** Dynamic organization of microtubules during oocyte development at early and mature Bb stages is concomitant with Bb condensation. Top: 2D images of early (left) and mature Bb (right) stage oocytes from Buc-GFP (green) ovaries labeled for microtubules (α-tubulin, inverted LUT). Zoom in of individual channel- and composite images are shown. Scale bars are 10 μm and 3 μm in the zoom out and zoomed in images, respectively. Bottom: 3D images of the cleft or mature Bb areas from the same oocytes above show that microtubules (magenta) are interwoven with the early (left) Buc condensate (cyan), but then encapsulate the mature Bb (right). XY, YZ and XZ images are shown. Scale bars are 2 μm. Images are snapshots form Videos S6. **B.** Transgenic EMTB-GFP (green) confirms microtubule organization during oocyte development at early and mature Bb stages, in fixed (top) and live (bottom) ovaries. DNA (blue) is labeled by DAPI. Scale bars are 10 μm. See Figure S3A. **C.** dSTORM super-resolution microscopy shows that Buc granules (red) are located immediately adjacent to microtubule cables (EMTB-GFP, green). Shown are representative Gaussian visualization images. i and ii are zoom-in images of color-coded boxes in the left panel. Scale bars are 100 nm. See Figure S3B. **D-O.** FRAP analyses of Buc-GFP turnover in the early forming Bb in the nuclear cleft, in whole ovaries treated with DMSO as control, compared to nocodazole (**D-G**), or colchicine (**H-K**), or taxol (**L-O**). Images (top DMOS-control, bottom drug treated ovaries) are snapshots at indicated timepoints (minutes) from representative Videos S7 (nocodazole), S8 (colchicine), and S9 (taxol). Recovery plots (E, I, M), as well as quantification by mobile fraction calculations (F, J, N), and by intensity at last 20 frames (G, K, O) are shown. All bars are Mean ± SD. Raw data are shown in Figure S4. See also Video S6-9, Figures S3-4.

α-tubulin labeling in Buc-GFP ovaries revealed microtubule cables and bundles through the cytoplasm at both cleft and mature Bb stages (Figure 4A). In early cleft stages, a prominent microtubule meshwork in the nuclear cleft co-localized with the Buc condensate (Figure 4A). In 3D views, microtubules appeared interwoven with the Buc condensate (n=21 oocytes, 6 ovaries; Figure 4A, Video S6). In mature Bb stages, microtubules formed a cage around the Bb (Figure 4A, Video S7). 3D views confirmed that the mature Bb appears to be encapsulated by microtubules in this cage-like organization (n=18 oocytes, 6 ovaries; Figure 4A, Video S6). EMTB-GFP ovaries (15 fixed, 20 live ovaries; Figure 4B) confirmed this organization, (n=15 fixed, 20 live ovaries) (Figure 4B), and Buc co-localization, confirmed it in respect to the cleft Buc condensate and the mature Bb (n=6 ovaries; Figure S3A).

The microtubule meshwork in the cleft aligns developmentally with early Buc condensation, positioning microtubules to regulate Buc dynamics. dSTORM microscopy confirmed the co-organization of Buc granules and microtubules (n=4 ovaries; Figures 4C, S3B). Buc granules were closely adjacent to microtubule cables in the cleft (n=4 ovaries; Figures 4C, S3B), suggesting microtubules may transport condensing Buc granules via a trafficking mechanism.

To test microtubule roles in Buc turnover, we performed FRAP on Buc-GFP ovaries treated with microtubule-depolymerizing drugs (nocodazole, colchicine) or DMSO as control (Figure 4D-K, S4, Videos S7-8). EMTB-GFP confirmed microtubule depolymerization in drug-treated ovaries, but not in DMSO controls (Figure S4G-H). In FRAP experiments, DMSO-treated oocytes showed ∼40-50% Buc-GFP recovery (n=17 oocytes; Figures 4D-K, S4, Videos S7-8), consistent with untreated controls (Figure 3). With nocodazole (n=20) or colchicine (n=14), recovery was reduced to ∼10-15% (Figures 4D-K, S4, Videos S7-8), confirmed by FRAP quantification (Figure 4D-K).

We repeated these experiments in Buc-GFP ovaries treated with the microtubule-stabilizing drug Taxol, confirming microtubules were not depolymerized (Figure S4I). Buc-GFP showed normal recovery in Taxol-treated ovaries (n=17; Figure 4L-O, Video S9), similar to DMSO controls (n=14). Recovery was slightly higher in Taxol-treated samples (Figure 4L-O). Mitotracker signals before and after FRAP ruled out cellular damage or artifacts from drug treatments (Figure S4J-L). We conclude that microtubules are necessary for Buc-GFP recovery and turnover in the early Bb condensate.

### Dynein-mediated trafficking of condensing Buc granules is required for Buc turnover

Buc granules localize towards the centrosome at the vicinity of the cleft^12^. Microtubules may transport Buc granules towards the centrosome via dynein. To test dynein’s role in Buc turnover, we conducted FRAP on Buc-GFP ovaries treated with DMSO or the dynein inhibitor ciliobrevin (Figures 5A-D, S5A-B, Video S10). Unlike DMSO-treated ovaries (n=20), ciliobrevin reduced Buc-GFP recovery to ∼10% (n=15; Figures 5A-D, S5A-B, F-H, Video S10), similar to microtubule depolymerization effects, indicating dynein’s necessity for Buc turnover in early Bb formation.

**Figure 5.**
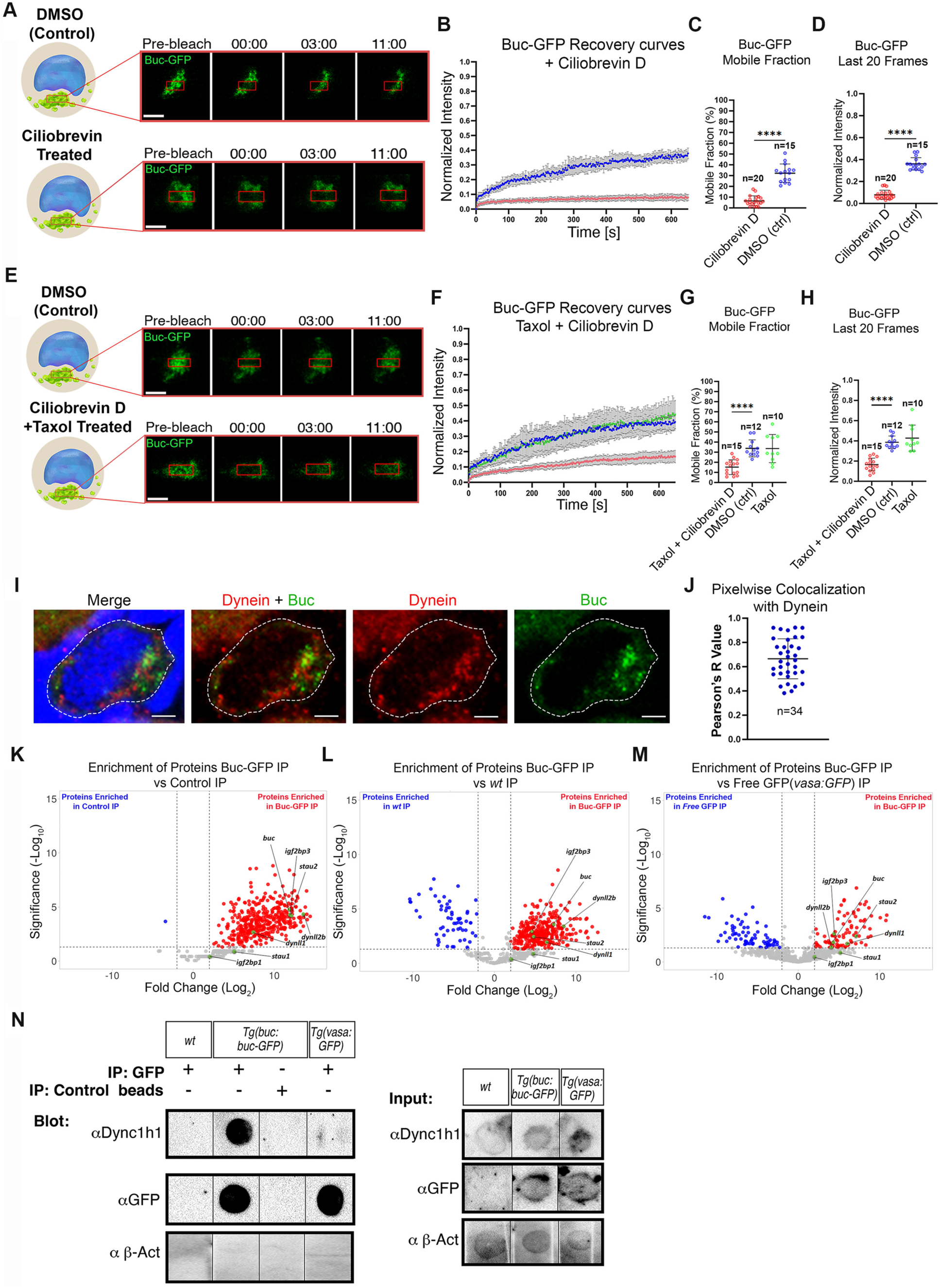
Early Buc turnover requires dynein-mediated microtubule trafficking. **A-H.** FRAP analyses of Buc-GFP turnover in the early forming Bb in the nuclear cleft, in whole ovaries treated with DMSO as control, compared to ciliobrevin (A-D), or to taxol and ciliobrevin (E-H). Images in A,E are snapshots at indicated timepoints (minutes) from representative Videos S10 (ciliobrevin) and S11 (taxol+ciliobrevin). Recovery plots (B, F), as well as quantification by mobile (C, G) fraction calculations, and by intensity at last 20 frames (D, H) are shown. All bars are Mean ± SD. Raw data is in Figure S5. **I.** Ovaries co-labeled for dynein (red), Buc (green), and DNA (DAPI, blue), show dynein localization to the nuclear cleft. Scale bar is 5 μm. **J.** Pixelwise colocalization test in oocyte nuclear cleft ROIs confirms dynein co-localization with Buc in the nuclear cleft. Pearson’s R value between 0 and 1 concludes co-localization. Bars are Mean ± SD. **K-M.** Buc-GFP-IP on oocytes isolated as in Figure 2A, followed by mass-spectrometry analysis. Volcano plots of proteins enriched specifically in Buc-GFP-IP, versus three independent controls: Control IP (no beads, M), no transgenic GFP IP (wt, N), and no Buc, free GFP-IP (vasa:GFP, O), showing specific pull down of dynein accessory subunits Dynll1 and Dynll2b, as well as known Bb proteins. Additional controls and raw data are shown in Figure S6. **N.** Dot-blot analysis of IP experiments as in M-O. IP (left) and input (right) samples are shown. Dynein heavy chain Dync1h1 is specifically pulled down only in Buc-GFP-IP samples, and it is detected in the input of all samples. GFP is detected in Buc-GFP and vasa:GFP IP and input samples, using anti-GFP beads, but not in control beads or wt (no GFP) IP and input samples. As specificity control, β-Act is not pulled down by Buc-GFP, and it is detected in all input samples. Raw blots are shown in Figure S7. See also Videos S10-11, Figures S5-7, Table S2.

Dynein can anchor microtubules to various cellular structures and thus may affect microtubule organization and/or stabilization^66^. To examine if dynein’s role is direct or via microtubule organization, we assessed microtubule structure after ciliobrevin treatment, finding it unchanged (Figure S5F), suggesting a direct trafficking role. We then repeated FRAP with Taxol pre-treatment to stabilize microtubules before ciliobrevin addition (Figures S5E-H, 5C-D, Video S11). If Buc-GFP turnover remains blocked despite stabilization, this supports a direct trafficking mechanism.

The organization of microtubules in DMSO control, non-treated, and Taxol plus ciliobrevin co-treated ovaries was indistinguishable (Figure S5G). FRAP experiments showed ∼40% recovery in DMSO (n=10) and Taxol-treated ovaries (n=12). However, Taxol plus ciliobrevin co-treatment (n=15) reduced Buc-GFP recovery to ∼15% (Figure S5E-H, S5D, Video S11), mirroring ciliobrevin alone and supporting a direct role of dynein in Buc trafficking. Supporting Buc trafficking, dynein co-localized with Buc in early cleft stage oocytes (Figure 5I), with puncta seen in the cytoplasm and cleft. Pixelwise colocalization analysis confirmed dynein-Buc co-localization in the cleft (Pearson’s R value=0.665±0.16; n=31 oocytes; Figure 5J).

We tested if Buc and dynein form protein complexes in developing oocytes during Bb formation. Using Buc-GFP ovaries, we performed immune-precipitation (IP) on isolated oocytes as in Figure 2 with Buc-GFP as bait, followed by mass spectrometry (Figure 5K-M, S6; Methods). Controls included: ***1)*** Buc-GFP IP versus control beads (Figures 5K, S6C), ***2)*** Buc-GFP IP versus WT ovaries without GFP (Figure 5L, S6C), and ***3)*** GFP IP of free GFP encoded protein from [*Tg(vasa:GFP)*]^67^ transgenic line (Figures 5O, S6C). IP specificity was confirmed by western blot and silver-stain (Figure S5A-B). Mass-spectrometry (MS) identified 91 proteins enriched in Buc-GFP IP compared to controls (Figures 5K-M, S6C-D, Table S2).

Enriched proteins included known Bb proteins (e.g., Igf2bp3), the Buc protein, and MARDO (mammalian oocytes RNA storage bodies)-associated proteins^68^ like Zar1, Ybx1, 4B-T, LSM14, and DDX6 (Figures 5K-M, S6C-D, Table S2). Gene ontology analysis showed many enriched proteins related to RNA processing and RNP granules (Figure S6E). Numerous proteins had IDRs and a PLD (Figure S6G), consistent with Buc-dependent insoluble Bb complexes (Figure 2). Dynein light chain subunits LC8-type 1 (Dynll) and LC8-type 2b (Dynll2b) were specifically enriched in Buc-GFP IP (Figures 5K-M, S6C-D). Those subunits act as dynein accessory components that link dynein to cargos and to adapter proteins that regulate dynein function^69^, supporting dynein transport of Buc.

Buc-dynein interaction was confirmed via Buc-GFP IP of dynein heavy chain Dync1h1, detected by dot blot (Figures 5N, S7). We confirmed specific IP of GFP in Buc-GFP and free GFP samples, but not in WT IP or control bead samples (Figures 5N, S7C). Dynein heavy chain was detected in all input samples but specifically in Buc-GFP IP and not in free GFP or control IPs (Figures 5N, S7). β-actin, used as a loading control, was absent in Buc-GFP IP with dynein heavy chain, confirming IP specificity (Figure 5N). These results support Buc binding to dynein subunits: two accessory light chains and the heavy chain.

In summary, three lines of evidence suggest dynein transports condensing Buc granules on microtubules during early Bb formation: ***1)*** Buc granules co-localize with dynein and microtubules in the cleft, ***2)*** Microtubules and dynein activity are essential for Buc-GFP turnover in the Bb condensate, and ***3)*** Buc forms complexes with Dync1h1, Dynll, and Dynll2b, likely representing a dynein transport complex.

### A microtubule meshwork scaffolds the main Buc condensate

To investigate if microtubules control already condensed granules after formation of the main condensate, we treated Buc-GFP ovaries with nocodazole, then fixed and co-stained for Buc-GFP, and two cleft landmarks: mAb414 (nuclear pore complexes) delineating the nuclear cleft (Figures 6A, S9), and γTub marking the centrosome in the vicinity of the cleft^12^ (Figure 6A). In DMSO-treated ovaries, 78.5±4.95% of oocytes had intact condensates in the cleft (n=126 oocytes, 6 ovaries; Figure 6A-B). Nocodazole-treated ovaries showed intact condensates in only 45.6±11.24%, with 54.3±11.24% exhibiting dispersed granules (n=163 oocytes, 6 ovaries; Figure 6A-B), indicating condensate disintegration upon microtubule depolymerization. We quantified these phenotypes by measuring Buc intensities along a defined line across the cleft (Methods), based on γTub (Figure 6D) or mAb414 landmarks (Figures 6D, S9A, E). Buc intensities were normalized and plotted along this line (Figure 6F-G; Methods). Figure 6E-J presents pooled data from two representative experiments, with individual raw data in Figure S8

**Figure 6.**
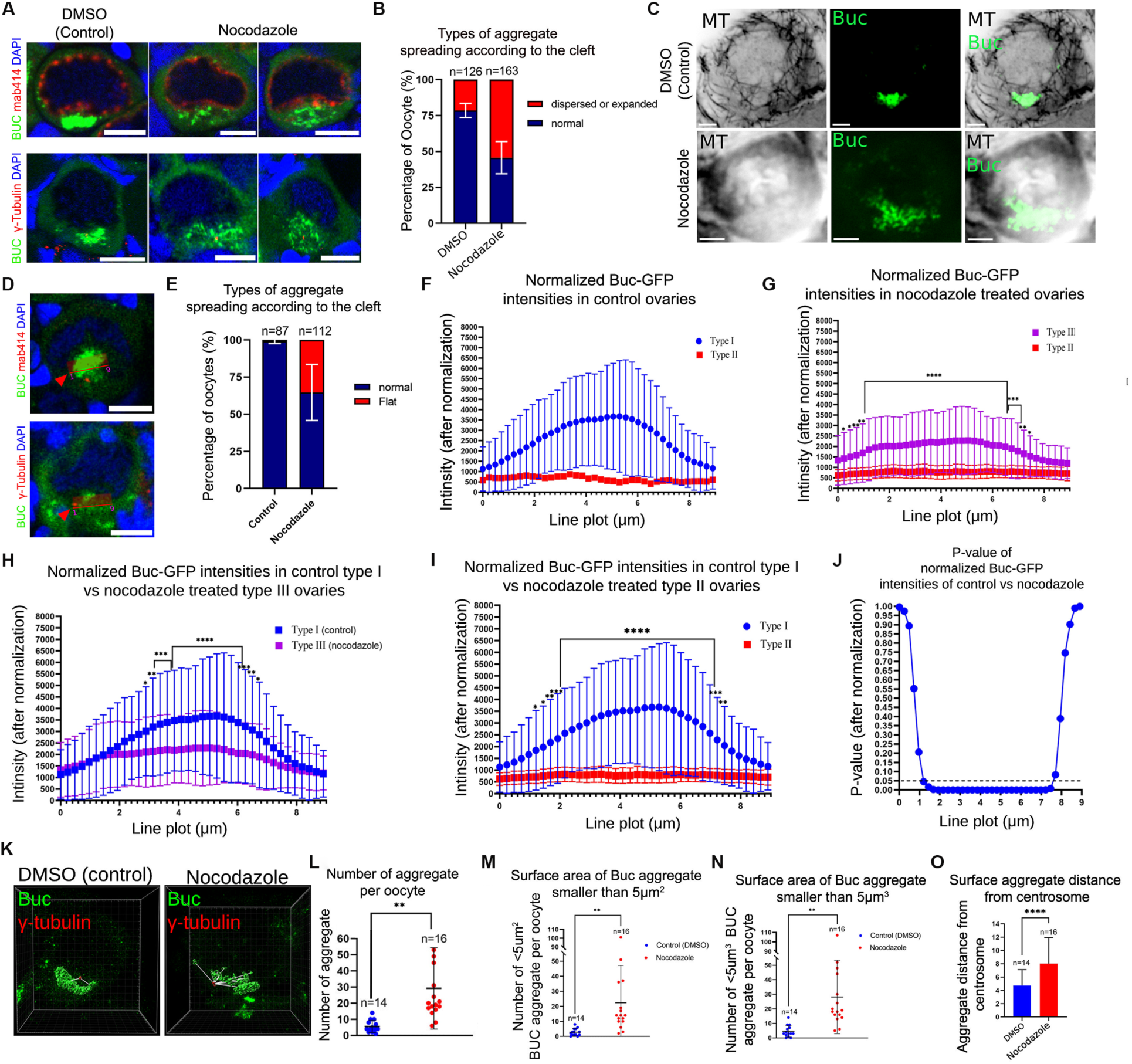
Microtubules are required for the integrity of the Buc condensate. **A.** Ovaries treated live with control DMSO (left) or nocodazole (right) and then fixed and co-labeled for mAb414 and Buc (top), or for γTub and Buc (bottom) show Buc condensate disintegration upon microtubule depolymerization. DNA (blue) is labeled by DAPI. **B.** Buc phenotype distribution from the experiments in A, are plotted. Bars are Mean ± SD from independent experiments. **C.** Representative images of EMTB-GFP ovaries co-labeled for EMTB-GFP (inverted LUT) and Buc (green), which were treated side by side with ovaries in A, showing microtubule depolymerization and Buc disintegration in nocodazole-treated, but DMSO-treated control ovaries. (n=6 ovaries for DMSO, and n=6 ovaries nocodazole). scale bars are 5 μm. **D.** Examples of the measuring line (arrowheads) of Buc intensity across the cleft and the forming Bb condensate, using the centrosome (γTub) as a marker for the cleft vicinity. Buc intensity plots were classified as type I (blue in E) or type II (red in E; Methods). DNA (blue) is labeled by DAPI. **E.** Plot of the rates of each curve types in control and nocodazole treated ovaries. **F.** Pooled Buc intensity curves in DMSO control ovaries. Type I exhibited a clear intensity peak, and type II were flattened with intensity <25% of maximal Type I peak. **G.** Pooled Buc intensity curves in nocodazole treated ovaries. **H.** Comparison of type I curves from control DMSO (blue) versus nocodazole treated ovaries (purple) show that type I curves in nocodazole samples exhibit significantly dampened peaks (asterixis indicate significant *p*-values). Thus nocodazole type I curves were re-classified as type III. **I.** Comparison of type I and type II curves show they are highly significantly distinct. **J.** Plot of the *p*-values of the comparison in I at each line position. **K.** Representative IMARIS 3D images of entire oocytes co-labeled for Buc (green) and γTub (red). White lines indicate the distance between automatically identified surfaces of Buc entities and the centrosome in control DMSO (left) or nocodazole (right) treated ovaries. Images are snapshots from Videos S12. **L-O.** The number per oocyte, of granules (L), number of granules with surface area smaller than 5 μm^2^ (M), number of granules with volume smaller than 5 μm^3^ (N), and the distance between granules and the centrosome (O), are plotted for control DMSO (blue) and nocodazole treated ovaries (red). All bars are Mean ± SD from independent experiments. See also Figures S8-9, Video S12.

In DMSO and nocodazole-treated ovaries, we observed two Buc intensity plot types. Type-I, with a clear peak, represents prominent Buc condensates (blue curves, Figures 6F-G, S8). Type-II, with intensities <25% of type-I maximal intensity and appearing flattened, likely indicates disintegrating condensates (red curves, Figures 6F, S8). In DMSO-treated ovaries, 99±1.41% of oocytes had type-I curves, with only 1±1.41% showing type-II (n=87, 6 ovaries; Figure 6E). In nocodazole-treated samples, 64.65±18.88% showed type-I, and 35.35±18.88% had type-II curves (n=112, 6 ovaries; Figure 6).

To assess microtubule depolymerization effects on Buc condensate integrity, we compared flattened type-II curves in nocodazole-treated oocytes (red, Figure 6G) with normal type-I curves in DMSO controls (blue, Figure 6F). Differences were statistically significant, with the lowest *p*-values near the center of the measuring line (Figure 6I-J), indicating Buc dispersal upon depolymerization.

Type-I-like curves in nocodazole-treated oocytes (purple, Figure 6G, H) were significantly dampened compared to type-I in controls (blue, Figure 6F, H), leading us to re-define them as type-III. This re-analysis showed that even oocytes previously considered normal with properly intact main condensates (Figure 6D, blue) had reduced Buc condensate intensity post-depolymerization, indicating dispersal and possibly dissolution of Buc. Results using γTub as a landmark (Figure 6, S8) were consistent with those using mAb414 (Figure S9)

We analyzed Buc condensate phenotypes in representative control oocytes with normal Bb condensate, and nocodazole treated oocytes with expanded and dispersed granules from the above experiments, in three dimensions. Control oocytes with a normal Bb condensate showed a prominent 3D Buc condensate adjacent to the centrosome (n=16; Figure 6K, Video S12). In nocodazole-treated oocytes, dispersed and expanded granules formed multiple smaller aggregates (n=14; Figure 6K, Videos S12). Microtubule depolymerization increased the total number of aggregates per oocyte, smaller aggregates per oocyte, and the aggregate distance to the centrosome (Figures 6L-O, S9K-L), indicating Buc condensate disintegration.

In conclusion, the microtubule meshwork is crucial for Buc condensate integrity in the cleft, potentially by localizing additional factors and/or by acting as a scaffolding molecular crowding agent.

### Microtubules encapsulate the mature Bb to prevent its over-growth and shape distortion

Later in development, microtubules form a cage-like structure around the mature Bb (Figure 4). To examine the role of this organization, isolated Buc-GFP oocytes were cultured with nocodazole or DMSO and monitored by time-lapse imaging (Methods). Microtubules in side-by-side EMTB-GFP oocytes depolymerized after ∼60 min (Figure S10A), marking T_0_ for 90-min Buc-GFP recordings (Methods). MitoTracker confirmed oocyte viability throughout.

In DMSO-treated oocytes, Bb size remained stable with slight fluctuations (Figure 7A-B, Video S13). Nocodazole-treated oocytes showed Bb inflation and growth of amorphic protrusions (Figure 7A-B, Video S13). Quantification (Methods) revealed a 2.26±0.44-fold increase in Bb volume and a 2.16±0.23-fold increase in surface area over 90 min (Figures 7C, S10B), with oocyte size unaffected in both treatments (Figure 7C). The Bb size increase could stem from accelerated Buc-GFP supply, but Bb density (Methods) decreased with growth to 69.66±7.09% of its initial density (Figures 7C, S10B), consistent with low Buc-GFP turnover (Figure 3), ruling out accelerated supply as a cause.

**Figure 7.**
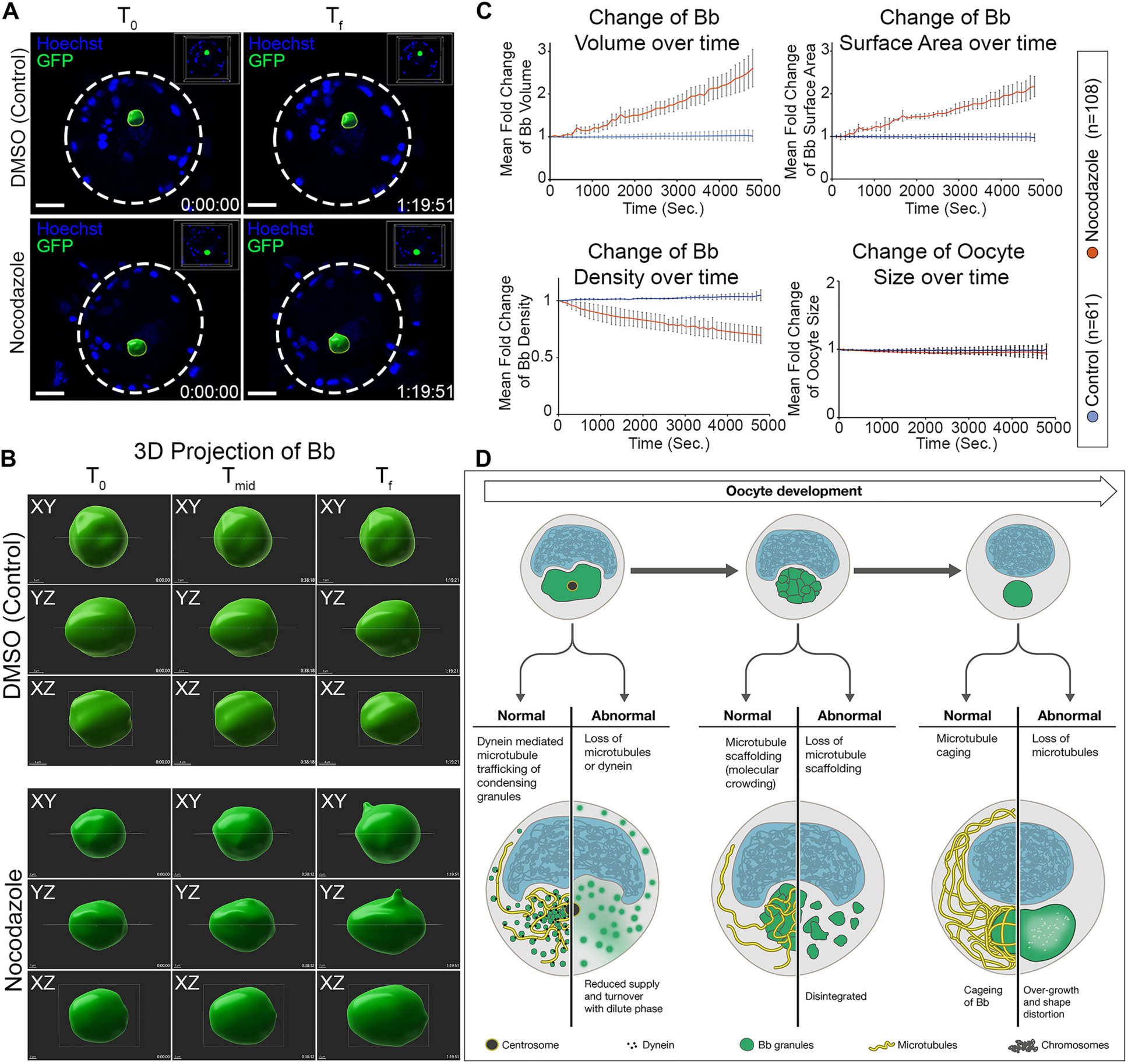
Microtubules restrict the over-growth and maintain the shape of the mature Bb condensate. **A.** Representative IMARIS 3D live-images of Buc-GFP control (top) and nocodazole treated (bottom) oocytes at T*_0_* (left) and T*_f_* (right). Oocyte contour is outlined by dashed white outline. DNA (blue) is labeled by Hoechst. Scale bars are 10 μm. **B.** Zoom in images of the Buc-GFP labeled Bb in control (top) and nocodazole (bottom) oocytes in the indicated projections and over time at three timepoints along the experiment. Images in A-B are snapshots of Video S13. **C.** Plots of fold-change over time in normalized Bb volume, surface area, and density, as well as of oocyte size, in DMSO control (blue) and nocodazole treated oocytes (orange). All bars are Mean ± SD from independent experiments. See Figure S10. **D.** Multistep control by microtubules of Buc condensation and Bb formation during oocyte development. Left: Early condensing Buc granules are trafficked by dynein on microtubules to the cleft, where they fuse and form the main early Bb condensate. The localization of Buc granules and Buc turnover in the condensate are microtubule and dynein dependent. Middle: At early-mid stages, microtubules scaffold the main condensate, and are required for its integrity, likely by providing molecular crowding agents to maintain Buc at concentrations that are sufficient for its condensed state or to support granule fusion. Right: At mature Bb stages, microtubule cage the Bb and prevents its overgrowth and shape distortion. See also Video S13, Figure S10.

These results show that microtubules are essential for maintaining Bb morphology and confining its size. Previous studies found no Bb degradation upon nocodazole-induced microtubule depolymerization^6,48^ but did not assess Bb dynamics or size increase over time. Genetic studies indicate Bb size must be regulated. Loss-of-function mutations in Macf1, which mediates cytoskeletal interactions, led to enlarged Bb, improper docking, radial oocytes, and infertility^6^, suggesting oversized Bb is non-functional.

## Discussion

Our work reveals mechanisms of Bb formation in vertebrates, addressing a question dating back to the Bb’s discovery in1845^9^. We show that Bb formation is driven by molecular condensation, as determined by the defining criteria of condensates^19,44,45,50^. First, early Buc IDP granules undergo density transitions, exhibit liquid-like properties, translocate and fuse, and exchange Buc with the dilute phase. Second, the condensate’s apparent viscoelastic properties transition over time to a solid-like state, showing reduced Buc exchange, resistance to IDR interference, and formation of presumptive amyloid structures dependent on Buc. Third, biochemical evidence confirms that the Bb contains Buc-dependent insoluble RNP complexes at the correct developmental stages. We conclude that Bb formation is driven by Buc phase-separation (and/or PS++), with presumptive viscoelastic properties that evolve developmentally (Figure 7D).

We identified mechanisms that regulate Bb condensation, from the initial localization of seeding Buc protein to the maturation of the compartment, through multi-step microtubule regulation (Figure 7D). First, dynein-mediated microtubule trafficking of early Buc granules is essential for Buc turnover in the forming condensate (Figure 7D, left). Second, a microtubule meshwork in the main condensate prevents disintegration into smaller granules and Buc dissolution (Figure 7D, middle). Finally, microtubules encase the mature Bb to control its growth and prevent shape distortion (Figure 7D, right).

These regulatory steps ensure a single, intact Bb condensate, which is considered crucial for oocyte polarization and development, and for embryogenesis. In fish, frogs, and most vertebrates, the Bb stores RNP complexes needed for maternal embryonic regulation, including key dorsal and germ cell determinants^1-3^. After delivery to the vegetal pole, essential RNP complexes remain at the vegetal pole in the developing embryo, from which they later guide axis formation^1-3^. The mechanisms we reveal provide insight into the origins of embryonic polarity in fish, frogs, and likely most non-mammalian vertebrates.

Our insights into the Bb provide a new perspective on oocyte biology across species, from insects to humans, where the Bb is evolutionarily conserved. In zebrafish, mice, and humans, the Bb follows similar developmental dynamics at corresponding oogenesis stages^9,12,13,15,70^. The process of oocyte symmetry-breaking, where Bb granules initially localize around the oocyte centrosome, is also conserved in insects, zebrafish, and mice^12,15,71^. In mice, a microtubule-dependent mechanism transports Bb organelles from nurse-like cells to the oocyte^72,73^, and Bb recruitment of Golgi elements depends on tubulins Tuba1b and Tubb2b^73^. Our findings emphasize molecular condensation as a critical factor in the developing vertebrate ovary, with shared Bb features suggesting conserved condensation mechanisms in mammals, relevant to human female reproduction.

Our work highlights the importance of cellular regulation over self-assembling phase-separation processes, as shown previously for P granules in *C. elegans*^74-77^ and pole granules in *Drosophila*^78-81,82,83^ embryos. We demonstrate similar microtubule-based control of oocyte polarity in zebrafish, suggesting this mechanism is conserved. In *Drosophila*, microtubules transport germ plasm condensates to (+)end tips via the kinesin motor^80,82^, and partition these condensates between daughter cells in *Drosophila* pole cells^83^ and zebrafish PGCs^84^ through the dynein motor. In polarized mammalian epithelia, microtubules are necessary for the apical localization of the Cobl protein, essential for the apical junction complex and likely forming condensates^85^. Increasingly, the relationship between molecular condensates and microtubules is recognized (rev. in^86,87^), underscoring the broad relevance of microtubule control over phase-separation.

Our findings offer insights into additional roles of microtubules in condensate regulation. *In-vitro* TIRF studies have shown that microtubules can drive condensation^85^, and (+)end enrichment in *Drosophila* oocytes was sufficient to assemble germ plasm^82^. Plausibly, the association of Buc complexes with dynein on microtubules may concentrate Buc locally, promoting condensation. Later, the cleft microtubule meshwork likely acts as a molecular crowding agent, preventing condensate disintegration and supporting fusion by maintaining Buc in a condensed state. *In-vitro*, microtubules can substitute for the crowding agent polyethylene glycol (PEG)^85^, suggesting a similar mechanism *in-vivo*.

Microtubules form a cage around the mature Bb, controlling its growth and preventing shape distortion. Bb overgrowth does not align with Ostwald ripening of condensates^64,88^, where the condensed phase expands at the expense of the dilute phase, retaining a spherical shape^64,88^. The reduction in Buc-GFP density within the enlarging Bb, combined with low Buc turnover, argues against Buc shifting from the dilute phase, and the growth of amorphic protrusion distorts the Bb spherical shape. Alternative mechanisms, suggested by theoretical and *in-vitro* models (e.g.,^89-91^), may apply. Mechanistically, microtubules could limit Bb growth by supplying or removing an unknown regulator or by directly forming a barrier. The cage-like microtubule structure encapsulating the mature Bb, along with the Bb’s solid-like properties and low cytoplasmic exchange, supports a direct role. Direct and indirect mechanisms may cooperate in regulating the mature Bb

Further understanding of Bb regulation is still needed. Previous studies on Bb transcript localization identified potent motifs in 3’UTRs of individual mRNAs (rev. in. ^8^), but a universal “zip code” motif has not been found. The discovery that Bb formation involves molecular condensation shifts focus from localization motifs to regions in proteins and RNA that act as stickers and spacers^32,33,44,45^, pointing investigation in the right direction. This can also address the still open question of the mechanisms that confer Bb constituents specificity.

The mechanism behind the Bb’s transition from liquid-like to solid-like states also remains unclear. This likely involves increasing saturation of Buc IDR interactions^92,93^ and possibly activity through Buc’s prion-like domain (PLD), which is likely functional^10^. In *Drosophila*, Bruno’s PLD is essential for hardening *oskar* RNA condensates^28^. Buc IDR interactions or PLD activity may be regulated by post-translational modifications; for example, Tdrd6a modulates Buc aggregation via methylation^94^. Buc is the only known essential Bb protein across species, with only two Buc-interacting proteins identified in oocytes: Tdrd6a^94^ and Rbpms2^95^, neither essential for Bb formation^94,95^. Asz1, another Bb protein in fish and frogs^12,96,97^, was recently shown not to be required for Bb formation^98^. Our updated list of Bb proteins interacting with Buc at specific developmental stages offers new candidates for future research. Overall, our work enhances understanding of Bb biology, oogenesis, and female reproduction.

## Resource Availability

### Lead Contact

Requests for further information and resources should be directed to and will be fulfilled by the lead contact, Yaniv M. Elkouby.

### Material Availability

This study did not generate new unique reagents.

### Data and Code Availability

All data, including proteomic data and original western blots are included with this manuscript. Microscopy data reported in this paper will be shared by the lead contact upon request.

Data reported in this paper will be shared by the lead contact upon request.

Any additional information required to reanalyze the data reported in this paper is available from the lead contact upon request.

## Acknowledgments

We are grateful to Roland Dosch, Rene Ketting, and Karuna Sampath for the *Tg (buc:Buc-GFP), Tg (vasa:GFP),* and *Tg (bact:EMTB-3XGFP)* lines. We thank Yaron Shav-Tal for his valuable advice on the analyses of our *in-vivo* FRAP experiments, and Reuven Vainer for his valuable advice on the Buc IP and supernatant/pellet experiments, Rohit Pappu and Eithan Lerner for their valuable advice on phase-separation, and Zakharya Manevitch for his help with SIM microscopy. We thank Mayyan Visuals for generating the cartoons in Figure 7D and the graphical abstract. This work was funded by the Israel Science Foundation – grant no. 558/19 to YME.

## Author contributions

Conceptualization: RD, SK, AA, YB, YME. Methodology: RD, SK, AA, YB, AD, GS, ES, AM, ABZ, YME. Investigation: RD, SK, AA, YB, AD, GS, ES, AM. Visualization: RD, SK, ES, YME. Funding acquisition: YME. Supervision: ABZ, YME. Writing – original draft: RD, SK, AA, YME

## STAR Methods

### EXPERIMENTAL MODEL

#### Fish lines

Fish lines used in this research are: Tü *wt*, *Tg (buc:buc-gfp-buc 3’utr*)^99^, *buc^p43^* ^100^ and *Tg (bact:EMTB-3XGFP)*^101^, *Tg (vasa:GFP)*^67^, *Tg(h2a:H2A-GFP)^102^*.

#### Ethics statement

All animal experiments were supervised by the Hebrew University Authority for Biological Models, according to IACUC and accredited by AAALAC. All experiments were appropriately approved under ethics requests MD-18-15600-2.

### METHOD DETAILS

#### Fish gonad collections

Juvenile ovaries were collected from 5-7 week post-fertilization (wpf) juvenile fish. Fish had a standard length (SL) measured according to^103^ and were consistently ∼10-15mm. Ovary collection was done as in^7,12^. Briefly, to fix the ovaries for immunostaining, fish were cut along the ventral midline and the lateral body wall was removed. The head and tail were removed and the trunk pieces, with the exposed abdomen containing the ovaries, were fixed in 4% PFA at 4°C overnight with nutation. Trunks were then washed in PBS and ovaries were finely dissected in cold PBS. Ovaries were washed in PBS and then either stored in PBS at 4°C in the dark, or dehydrated and stored in 100% MeOH at -20°C in the dark.

#### Genotyping

Genotyping was performed at 4 wpf and over maximum of 2 days, after which fish were rested and raised in the system until the collection of their ovaries at 5-7 wpf. Fish were anaesthetized in 0.02% Tricaine in system water (Sigma Aldrich, #A5040), and a piece of their fin tail was clipped for DNA extraction. DNA was extracted using the standard *HotSHOT* DNA preparation method. *Tg (buc:Buc-GFP)* and *Tg (vasa:GFP)* fish were genotyped by genomic PCR, using PCRBIO HS Taq Mix Red (PCR Biosystems #PB10.13-02), and primers targeting the *gfp* sequence: forward: GACGTAAACGGCCACAAGTTCAG, reverse: GTTCACCTTGATGCCGTTCTTC. *buc^p43^* mutant fish were genotyped by KASP genotyping (LGC, Teddington, UK). KASP-PCR was performed on an Applied Biosystems StepOnePlus machine and following the manufacturer instructions, and the SNP primer sequence: ATCCACTCTGTTGCCCCCACAAAACTATTTGTCCTCAATTGGCAGTGCTTATTCCTAC AGCTA[C/A]TACCCACAAGTGACCCAAGAGCGCCAGAGTGTCTTAAGTCCATCCATAGATGAGCTCTCC.

#### Fluorescence immunohistochemistry (IHC) and HCR-FISH

Fluorescence immunohistochemistry (IHC) was performed as in^7,12^. Briefly, ovaries were washed 2 times for 5 minutes (2x5min) in PBT (0.3% Triton X-100 in 1xPBS; if stored in MeOH, ovaries were gradually rehydrated first), then washed 4x20min in PBT. Ovaries were blocked for 1.5-2hrs in blocking solution (10% FBS in PBT) at room temperature, and then incubated with primary antibodies in blocking solution at 4°C overnight. Ovaries were washed 4x20min in PBT and incubated with secondary antibodies in fresh blocking solution for 1.75hr, and were light protected from this step onward. Ovaries were washed 4x20min in PBT and then incubated in PBT containing DAPI (1:1000, Molecular Probes), with or without DiOC6 (1:5000, Molecular Probes) for 50min and washed 2x5min in PBT and 2x5min in PBS. All steps were carried out with nutation. Ovaries were transferred into Vectashield (with DAPI, Vector Labs). Ovaries were finally mounted between two #1.5 coverslips using a 120μm spacer (Molecular Probes).

Primary antibodies used were mouse anti-GFP (1:300; Molecular Probes), rabbit anti-GFP (1:400; Molecular Probes) and rabbit anti-Buc^6^ (1:500; Yenzym Antibodies LLC), αTubulin (1:1000, Merck), γTubulin (1:400, Sigma-Aldrich), mAb414 (1:1000, Abcam), dynein heavy chain DYNC1H1^104^ (1:1000; Proteintek). Secondary antibodies used were Alexa Fluor 488 and 594 (1:500; Molecular Probes). Vital dyes used were: DiOC6 (1:5000, Molecular Probes)^7,12^, Mitotracker (1:2000, Molecular Probes).

RNA-FISH for *dazl* mRNA was performed using the DNA-HCR-FISH technique (Molecular Instruments)^105^, as in^7,12^ and following the company protocol, except for the hybridization temperature that was optimized for 45°C.

#### Confocal microscopy

Images were acquired on a Zeiss LSM 880 confocal microscope using a 40X lens. The acquisition setting was set between samples and experiments to: XY resolution=1104x1104 pixels, 12-bit, 2x sampling averaging, pixel dwell time=0.59sec, zoom=0.8X, pinhole adjusted to 1.1μm of Z thickness, increments between images in stacks were 0.53μm, laser power and gain were set in an antibody-dependent manner to 4-9% and 400-650, respectively, and below saturation condition. Unless otherwise noted, shown images are partial Sum Z-projection. Acquired images were not manipulated in a non-linear manner, and only contrast/brightness were adjusted. All Figures were made using Adobe Photoshop CC 2014.

#### Confocal live and time-lapse imaging of cultured ovaries

Confocal live and time-lapse imaging of cultured ovaries was performed as in^7,106^. Briefly, ovaries were dissected from juvenile fish (5-7wpf, SL∼10-15mm) into fresh warm (28°C) HL-15 media (Hanks solution containing 60% L-15 (no phenol red), Sigma-Aldrich, and 1:100 Glutamax, Invitrogen, warmed to 28°C). Ovaries were then embedded in 0.5% low-melt agarose in HL-15 on a glass-bottom dish, and covered with warm HL-15. After the agarose polymerized, ovaries were incubated in HL-15 at 28°C. Time-lapse images of live ovaries were acquired using either a Zeiss LSM 880 or Nikon Ti2E spinning disc confocal microscopes, both equipped with an environmental chamber set to 28°C, and using a 40X and 60X objectives, respectively. Images were acquired every 6 seconds for single Z section, or 12 seconds for Z stacks, over 5-10min.

#### SIM Super-resolution microscopy

Juvenile *Tg(buc:buc-GFP)* ovaries were dissected and fixed as described above. Fixed ovaries were washed X3 times in 1xPBS and mounted in ProLong Gold antifade reagent (Invitrogen) on Superfrost Plus Adhesion Microscope Slides (dimension: 25x75x1 mm) and covered with 18x18 mm High-Performance cover glasses (Zeiss). Slides were cured at 4°C overnight. Images were acquired in Nikon Ti2 Eclipse SIM microscope using a 100X TIRF objective, at XY=100 nm and Z=200 nm resolution. Images were mildly adjusted for contrast/brightness (with no non-linear manipulations) in Fiji, and 3D images were created by the designated Nikon NIS element software.

#### dSTORM Super-resolution microscopy

For super-resolution microscopy, we used a dSTORM system and protocol as described in^107^, which allows for imaging at approximately 20 nm resolution. All dSTORM imaging was performed on TIRF mode. Whole *Tg (βact:EMTB-3XGFP)* juvenile ovaries were immune-stained, using primary mouse anti-GFP (1:500; Molecular Probes) and rabbit anti-Buc (1:500; polyclonal antibodies against N-term, Buc peptide Yenzym Antibodies LLC) antibodies. Secondary antibodies used were anti-rabbit H+L Alexa Fluor 568 and anti-mouse H+L Alexa Fluor 488 (1:500; Life Technologies). After IHC, ovaries were then mounted on poly-D-lysine coated coverslips (no. 1.5H, Marienfeld-superior, Lauda-Königshofen, Germany). dSTORM imaging was performed in a freshly prepared imaging buffer containing 50 mM Tris (pH 8.0), 10 mM NaCl and 10% (w/v) glucose with an oxygen-scavenging GLOX solution (0.5 mg/ml glucose oxidase (Sigma-Aldrich), 40 µg/ml catalase (Sigma-Aldrich), 10 mM cysteamine MEA (Sigma-Aldrich) and 1% β-mercaptoethanol. A Nikon Ti-E inverted microscope was used. The N-STORM Nikon system was built on TIRF illumination using a 1.49 NA 100X oil immersion objective and an ANDOR DU-897 camera. 488, 568 nm laser lines were used for activation with cycle repeat of ∼4000 cycles for each channel. Nikon NIS Element software was used for acquisition and analysis. A gaussian visualization is shown in images in Figure 1D in which signal intensity correlates with localization precision. ‘Cross-visualization’ in Figure S3B shows all resolved signals where each single-molecule signal displays as a cross in the N-STORM software.

#### dSTORM quantification by Cluster analysis

Clustering quantification were performed as described previously *(107)*. In brief, *cluster signal densities*: Single molecule localization microscopy (SMLM) results in point patterns having specific coordinates of individual detected molecules. These coordinates are typically summarized in a ‘molecular list’ (provided by ThunderSTORM analysis (NIH ImageJ)). In order to define molecular clusters, we analyzed the molecular lists through a custom Matlab code (MathWorks) using the Matlab functions "Cluster" and "Linkage", as follows: First, our code calculated distances between each point and all other points in the point pattern of the SMLM image. Then, we set a distance threshold for defining molecules that belong to the same cluster: two points were defined to be clustered if their distance was smaller than the threshold distance (e.g. 70 nm). All points that were clustered with a specific point belong to one cluster (as defined by linkage function). Hence, a point could only be within one cluster. The code then defines and saves the properties of each cluster, such as the area of the cluster, the number of points within the cluster, and the number of clusters. Finally, the point patterns were visualized, while showing all points that belong to the same cluster with the same identifying color (Simulations in Figure 1D).

#### Fluorescence Recovery After Photobleaching (FRAP)

FRAP was performed on a Zeiss LSM 880 confocal microscope in ovaries carrying Buc-GFP transgene [*Tg(buc:buc-gfp-buc 3’utr*]. In preparation experiments, we established that bleaching 1/3 of the aggregate resulted in a significant reduction in Buc-GFP signal without completely ablating all the protein in the nuclear cleft. Furthermore, we observed a plateau of recovery dynamics after 680sec, and therefore to reduce any photobleaching affects, all experiments had a total duration of 680sec. Based on these calibration experiments, in all experiments we systemically bleached an area of approximately 1/3 of the aggregate size to around 20% of the original Buc-GFP intensity and then followed recovery for 680 seconds (11.33min), recording every 2.5 second. Regions (pre– and post-bleach) were tracked using ROI manager on Fiji. 10 pre-bleach frames were recorded and after background subtraction, the average intensity was used as pre-bleach intensity. Post-bleach frames were background subtracted and to make replicates comparable, post-bleach frame #1 of each measurement was set to 0 and corresponding pre-bleach intensity was corrected for this. Subsequently, recovery curves were plotted and the immobile/mobile fractions were calculated (see below) in order to determine Buc protein dynamics and turnover within individual oocytes. Mobile and Immobile fractions were calculated according to European Advanced Light Microscopy Network (EAMNET):

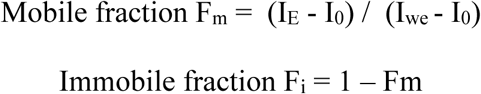

I_E_ is the end value of the recovered fluorescence intensity, I_0_ is the first postbleach fluorescence intensity, and I_I_ is the initial (prebleach) fluorescence intensity.

As control, FRAP was performed on ovaries, which transgenically express free GFP [*Tg(vasa:GFP)*], or Histon2A-GFP [*Tg(h2a:H2A-GFP)*], using the same settings as above. Free GFP recovery is very fast and requires minor adjustments of the size of bleached area, as well as recording speed^59^. Thus, to capture GFP bleaching, a smaller region was bleached and the entire experiment was recorded faster with 0.6 seconds intervals for 2.37 minutes, as previously suggested^59^. GFP recovery reached plateau at ∼2.4 seconds and continued to fluctuate thereafter because of the high dynamics and movement of the protein.

#### Thioflavin (ThT)

Ovaries were dissected from wt, *Tg (buc:buc-gfp-buc 3’utr*), or *buc^-/-^* juvenile fish (5-7 wpf, SL ∼10-15mm) into fresh warm (28°C) HL-15 media as described above and in (Elkouby & Mullins, 2017b). Ovaries were incubated with 20μM ThT (Sigma Aldrich #T3516) for 30 minutes and then rinsed x2 in fresh warm HL-15 media before being mounted for live imaging or fixed for staining.

#### Drug Treatments

Ovaries were dissected from wt, *Tg (buc:buc-gfp-buc 3’utr*), or *Tg (bact:EMTB-3XGFP)* juvenile fish (5-7 wpf, SL ∼10-15mm) into fresh warm (28°C) HL-15 media as described in^108-110^. Then HL-15 was replaced with HL-15 containing the drug of interest. For microtubule depolymerization and dynein inhibition, we used either 20μM nocodazole (Abcam #ab120630), 50μM colchicine (Sigma Aldrich #C9754), 40μM taxol (Abcam #ab120143), 25μM ciliobrevin (Sigma Aldrich #250401), or an equivalent volume of DMSO as vehicle control. Each drug treatment experiment contained a DMSO control group and for microtubule manipulating drugs also a *Tg (bact:EMTB-3XGFP)* sample as readout of the effects on microtubule. Ovaries were incubated for 90min at 28°C in the dark. Ovaries were then mounted for live imaging in media containing their respective drug concentration or DMSO control. Under these conditions, ovaries are normally viable for at least 8hrs^7^, and their viability was verified by MitoTracker Mitochondria-Selective Probes (500nM, Molecular Probes #M7512). At the end of treatments, ovaries were either continued to be imaged live, or fixed for analyses. For 1,6-hexanediol, live ovaries were similarly dissected. Ovaries were incubated in either 5% or 10% 1,6-hexanediol (Sigma, #240117) or equivalent volumes of MeOH as vehicle control, and either imaged live for 90 minutes while recording full z-stacks every 10 minutes, or treated for 90 minutes and then fixed for *dazl* HCR-FISH.

#### Stage Specific Oocyte Isolation

Ovaries were collected from adult females at 6-8 months post-fertilization. Stage specific oocyte isolation from ovaries was performed as described in^7,54^. Briefly, oocytes are first isolated from ovaries by short enzymatic digestion (3 mg/ml Collagenase I, Sigma-Aldrich, #C0130; 3 mg/ml Collagenase II, Sigma-Aldrich, #C6885; 1.6 mg/ml Hyaluronidase, in HL-15, Sigma-Aldrich, #H4272), at 28°C and in HL-15. Isolated oocytes are separated by size using cell strainers in various pore sizes (BD Falcon), and washed in HL-15. Oocytes are then immediately lysed for biochemical experiments, or cultured in HL-15 at 28°C for live experiments.

#### Oocyte lysis and separation of pellet and supernatant fractions

Stage specific oocytes were isolated as described above and were homogenized with a micro-pestle on ice in lysis buffer [10mM Tris (pH 7.5), 150mM NaCl, 0.5mM EDTA, 0.5% NP-40, 1X complete protease inhibitor cocktail (Roche, Mannheim)] according to (Riemer et al., 2015). After lysis, samples were spun down 10min at 12,000 x g at 4°C and the subsequent supernatant and pellet fractions were isolated separately for western blot and/or mass-spectrometry analyses. In Figure 2B, L-Arginine (Sigma Aldrich #5131) was added directly to the lysis buffer, prior to starting the western blot protocol.

#### Immunoprecipitation

Each immunoprecipitation sample was performed on oocytes isolated from 4 whole ovaries, which were dissected from 2 fish, as described above. Samples were lysed in 650 mL lysis/IP buffer [10mM Tris (pH 7.5), 150mM NaCl, 0.5mM EDTA, 0.5% NP-40, 1X complete protease inhibitor cocktail (Roche, Mannheim)] according to (Riemer et al., 2015) and homogenized with a micro-pestle followed by sonication for 3 x 30sec at low power. 25μl of GFP-Trap beads (GFP-Trap Dynabeads, Chromotek #gtd-20) per lysate were used for GFP pulldown from experiment *Tg (buc:Buc-GFP)*, as well as *wt* no transgenic GFP control, and *Tg (vasa:GFP)* free GFP control samples, and 25μl of control no-GFP nanobody beads (Chromotek), were used for IP control on *Tg (buc:Buc-GFP)* samples. Samples were then incubated for 1hr at 4°C, followed by 3 washes with wash buffer before western blot, mass-spectrometry, or dot blot analyses. Each sample (Buc-GFP-IP, and the three controls: no GFP-IP, beads only-IP, and free-GFP-IP), were performed in biological triplicates and the entire experiment was performed at least three times.

#### Mass-Spectrometry Analysis

All MS samples were performed and analyzed at the Smoler Proteomics Centre at the Technion (Haifa, Israel). The samples were precipitated in 80% acetone, and washed 4 times with cold 80% acetone. The protein pellets were dissolved in 8.5M Urea and 400mM ammonium bicarbonate and 10mM DTT. The samples were reduced (60°C for 30min), modified with 35.2mM iodoacetamide in 100mM ammonium bicarbonate (RT for 30min in the dark) and digested in 1.5M Urea, 66mM ammonium bicarbonate with modified trypsin (Promega), overnight at 37°C in a 1:50 (M/M) enzyme-to-substrate ratio. An additional second trypsinization was done for 4hrs. The resulting tryptic peptides were desalted using C18 tips (Home-made stage tips), dried and re-suspended in 0.1% Formic acid. They were analyzed by LC-MS/MS using Q Exactive HFX mass spectrometer (Thermo) fitted with a capillary HPLC (easy nLC 1000, Thermo). The peptides were loaded onto a homemade capillary column (20 cm, 75micron ID) packed with Reprosil C18-Aqua (Dr. Maisch GmbH, Germany) in solvent A (0.1% formic acid in water). The peptides mixture was resolved with a (5 to 28%) linear gradient of solvent B (95% acetonitrile with 0.1% formic acid) for 120min followed by gradient of 15min gradient of 28 to 95% and 15min at 95% acetonitrile with 0.1% formic acid in water at flow rates of 0.15μl/min. Mass spectrometry was performed in a positive mode using repetitively full MS scan followed by high collision induces dissociation (HCD, at 25 normalized collision energy) of the 10 most dominant ions (>1 charges) selected from the first MS scan. The mass spectrometry data was analyzed using the MaxQuant software 1.5.2.8. (//www.maxquant.org) using the Andromeda search engine, searching against the *Danio rerio* proteome from the Uniprot database with mass tolerance of 4.5ppm for the precursor masses and 4.5ppm for the fragment ions. Peptide- and protein-level false discovery rates (FDRs) were filtered to 1% using the target-decoy strategy. Protein table was filtered to eliminate the identifications from the reverse database, common contaminants and single peptide identifications. The data was quantified by label free analysis using the same software, based on extracted ion currents (XICs) of peptides enabling quantitation from each LC/MS run for each peptide identified in any of experiments. Statistical analysis of the identification and quantization results was done using Perseus 1.6.7.0 software.

#### Bioinformatical Analysis

Primary Statistical analysis was performed using Perseus 1.6.7.0 software. Differential expression of proteins has been calculated manually using Excel. For Sup/Pel and Buc-IP experiment, LFQ intensities of identified proteins for all samples were log normalized. To identify differentially enriched proteins, pairwise comparison was performed between experiment and control samples, using the formula *log2 (Fold Change) = log2 (protein intensity of sample1 / protein intensity of sample2)*, *p-*values were calculated by students t-test between samples. For Sup/Pel experiment pairwise comparison of fold change and *p*-value were calculated between *wt* sup Vs. *wt* pellet and *buc^-/-^* sup Vs. *buc^-/-^* pellet. For Buc-IP experiments, pairwise comparison of fold change and *p*-value were calculated between the Buc-GFP Vs. all other control samples. The cutoff for fold change has been set as 2 and for *p*-value<0.05. Enriched proteins for both sup/pel and BUC-IP experiments were visualized by Volcano Plots using the R-based software VolcanoseR^111^. Heatmaps and hierarchical clustering of enriched proteins ere generated using R and the R-based software ClustVis^112^.

#### Gene Ontology Analysis

Gene ontology analysis of enriched Bb resident and identified Buc binding proteins was performed using the software ShinyGo^113^ gene ontology database. The cut off for the FDR has been set as FDR<0.05.

#### Computational analyses of IDR and PLD

Primary sequences of enriched Bb resident and Buc binding proteins were taken from UniPort^114^ as FASTA sequence format and analyzed using three different algorithms, PONDR^115^, FoldIndex^116^ and ESpritz^117^, independently to predict protein ordered and disordered regions. PLAAC^118^ software was used to identify sequences of prion like domains in those proteins.

#### Western Blot analysis

Samples were heated to 95°C with 2X sample buffer (4X stock contains: 40% glycerol, 0.2M Tris pH 6.8, 20% ß-mercaptanol, 4% SDS and 0.005% Bromophenol Blue) for 5min prior to loading on a 4%-20% Mini-PROTEAN TGX gels (BioRad, #456-1094) and blotted using Trans-Blot Turbo Transfer Pack 0.2μm nitrocellulose (BioRad, #1704159) on the TransBlot Turbo BioRad system at 2.5A, 23V for 10min. Membranes were blocked for 2hrs at RT in 5% milk powder and then incubated overnight (o/n) at 4°C with primary rabbit anti-GFP antibody (1:1,000; Molecular Probes). Membranes were washed 3x for 10min in 1X TBST. Membranes were then incubated with the secondary antibody, peroxidase-conjugated AffiniPure Goat Anti-Rabbit IgG (H+L) (Jackson ImmunoResearch, #111-035-003), for 1hr at RT. Membranes were finally washed 5x for 5min in 1XTBST before being developed with the ECL Clarity Kit (BioRad) and imaged on ChemiDoc Imaging Systems (BioRad). Images were analyzed on Image Lab 4.1 (BioRad).

#### Dot Blot analysis

After Immunoprecipitation, samples were heated to 95°C with 2X sample buffer for 5 mins before loading the samples directly to a 0.2μm nitrocellulose (BioRad, #1704159) membrane. Membranes were cut as thin strips and grid lines have been drawn with pencil on the membrane. 10μl of each sample were directly loaded on each grid on the membrane and the membrane was air dried. Membranes were then blocked for 1 hour at RT in 5% milk powder and then incubated overnight (o/n) at 4°C with primary antibodies. Antibodies used were rabbit anti-GFP antibody (1:1,000; Molecular Probes), rabbit anti-DYNC1H1 antibody (1:1000; Proteintek), and mouse anti-b-Actin antibody (Jackson ImmunoResearch, #111-035-003). Membranes were washed 3x for 10min in 1X TBST and incubated with the secondary antibody, peroxidase-conjugated AffiniPure Goat Anti-Rabbit IgG (H+L) (Jackson ImmunoResearch, #111-035-003), or peroxidase-conjugated AffiniPure Goat Anti-Mouse IgG (H+L) (Jackson ImmunoResearch, #115-035-003) for 2hr at RT. Membrane strips were finally washed 6x for 3mins in 1X TBST before being developed with the ECL Clarity Kit (BioRad) and imaged on ChemiDoc Imaging system (BioRad). Images were analyzed on Image Lab 4.1 (BioRad).

#### Measurements of Buc intensity plots in the cleft

To analyze Buc condensates in the forming Bb, we analyzed the intensity and distribution of Buc-GFP signals in the cleft. The cleft was identified by antibody labeling detecting two landmarks: *1)* mAb414, detecting FG repeats in nuclear pore complexes and delineating the nuclear envelop (NE), and *2)* γTub, detecting the centrosome at the vicinity of the cleft, where Buc localizes. We identified the optical z-section in which the cleft was most pronounced based on the NE mAb414 signal, or the one that included the centrosome, both indicating the approximate center of the cleft and the main Buc condensate. We generated max z-projection images of three optical sections, including the one of the cleft center as identified above, as well as one below and one above it. We measured Buc intensities and distribution in the cleft, using the Fiji software. We drew a 9 μm line between the nuclear protrusions that encapsulated the cleft, and across the Buc condensate. Based on analyses on oocytes at these stages in preparation experiments, we determined that a line of 9 μm in length consistently spans the entire cleft opening and covers the main Buc condensate. The measuring line was divided into 37 bins every 0.24 µm and the intensity of Buc-GFP was measured at each bin. Buc-GFP signals at each bin were first background subtracted and then normalized. For measuring Buc-GFP background, we measured its intensity in a square (20 µm^2^) drawn in the oocyte nucleus, where Buc should not localize. To control for immune-labeling variation, background subtracted Buc-GFP signals were normalized to the mean intensity/area of co-labeled mab414 or γTub signals. Normalized Buc-GFP intensity were plotted along the measuring line for each oocyte. Pooled data was plotted as mean ± SD for each experiment and between experiments.

#### Three-dimensional analysis of Buc in IMARIS

Confocal raw data images of entire z-stacks of oocytes were extracted from whole ovary images and imported to IMARIS. Nuclei, centrosomes and Buc condensates were segmented in 3D using blend volume rendering mode, and signal brightness and object transparency were slightly and linearly adjusted in all channels to optimize visualization. Animation frames were made using the key frame animation tool and surfaces of centrosomes and Buc-GFP granules were generated. The threshold absolute intensity of Buc-GFP was set as 0.22 of Buc-GFP maximum intensity and the threshold of voxels was set as 0.003 of the maximum voxel number, for all oocytes across samples and experiments. To analyze Buc-GFP granules, their automatically generated number and surface area were extracted from IMARIS. To analyze the distance between Buc granules and the centrosome, the distance between the center of the mass of the centrosome and the center of mass of each Buc granule was measured in IMARIS. Measured parameters were exported to Excel and GraphPad Prism for statistical analyses and plotting.

#### Live Bb over-growth experiments and IMARIS analyses

Ovaries were dissected from *Tg (buc:buc-gfp-buc3’utr*), and *Tg (bact:EMTB-3XGFP)* adult fish into fresh warm HL-15 media as described in^7^. Ovaries were digested and oocytes at sizes of 50-100 μm were isolated as described above in the stage specific oocyte isolation section. After isolation, oocytes were embedded in low melt agarose with 20 μM nocodazole or DMSO in glass bottom dish and the dish was filled with HL-15 medium containing 20 μM nocodazole or DMSO, respectively. Oocytes from *EMTB-3XGFP* and *Buc-GFP* ovaries were incubated at 28°C in the dark and treated side by side, and the *EMTB-3XGFP* oocytes were imaged to monitor microtubules depolymerization. The time point of microtubule depolymerization (after ∼1 hour), as detected in the *EMTB-3XGFP* sample, was set as *T_0_* for Buc-GFP oocytes, which were then imaged for 1.5 hours, while recording z-stacks every 1 min. Live imaging was performed on a spinning disc confocal microscope. Live time-lapse images were exported to IMARIS for further analysis.

In the IMARIS software, each oocytes and their respective Bbs were segmented throughout the Z axis, and the *1)* volume, *2)* surface area and *3)* the intensity of Buc-GFP were recorded throughout the course of the 1.5 hours of imaging. Next, the absolute volume (*V_Bb_*), Surface area (*SA_Bb_*) and the Intensity (*I_Bb_*) of Bbs, as well as the volume (*V_O_*), Surface area (*SA_O_*) and the Intensity (*I_O_*) of their corresponding oocytes were recorded over time. The absolute volume, surface area and intensity were normalized to the oocyte volume, surface area and the intensity respectively. The density of the Bb was calculated using the normalized intensity of Bbs over their normalized volume, as follows:

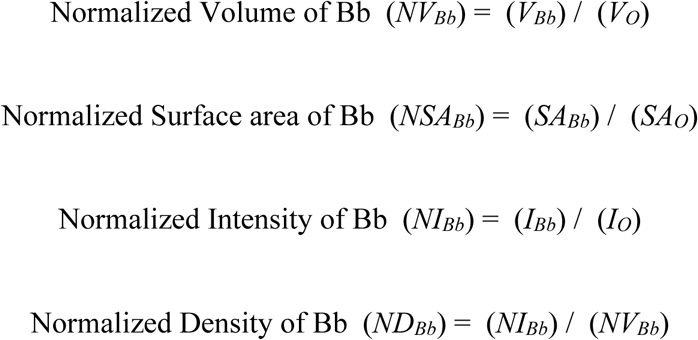

Next, the Fold change of the normalized volume, surface area and the density were calculated by dividing these normalized parameters at each time point (*T_i_*) to these normalized parameters at the starting time point (*T_0_*), as follows:

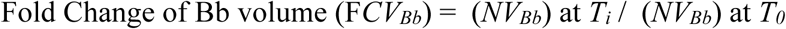

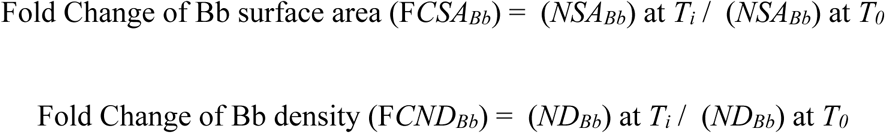

Finally, the fold change values of these parameters were plotted as the mean ±SD, over time.

### QUANTIFICATION AND STATISTICAL ANALYSIS

All statistical analysis and data plotting were performed using the GraphPad Prism 7 software and Microsoft Excel. Data sets were tested with two-tailed unpaired t-test. *p*-values were: *<0.01,****<0.05).

## Supplementary Information

**Document S1.** Figures S1-10.

**Table S1. Buc dependent Bb proteins. Related to Figure 2**. A list of the 114 Buc dependent enriched protein in the pellet over supernatant samples.

**Table S2. Buc interacting proteins. Related to Figure 4**. A list of the 90 enriched proteins specifically in Buc-IP samples.

**Video S1. 3D SIM of the early Buc condensate. Related to Figure 1B**. 3D SIM super-resolution views of early Bb condensate (Buc-GFP, green) in the cleft.

**Video S2. Live time-lapse imaging of Buc granules. Related to Figure 1E**. Live time-lapse imaging of Buc-GFP granules (green) in ovaries exhibit liquid droplet-like dynamics. The video shows an example of granule condensation (color-coded arrowheads tracking individual granule), followed by an example of granules that fuse with other granules (arrowheads), and finally an example of granules that translocate to and fuse with the main aggregates (arrowheads). XY and YZ views are shown. Scale bars are 2 μm. Time (hh:mm:ss) is indicated. We occasionally observed granule splitting; fusion and splitting dynamics are characteristics of liquid droplet-like condensates.

**Video S3. FRAP analyses of Buc in the early condensate versus the mature Bb. Related to Figure 3**. Live time-lapse imaging of FRAP analyses of Buc-GFP turnover in in whole ovaries, in the early forming Bb in the nuclear cleft (left) and in the mature Bb (right). White outline indicates oocyte contour, red boxes indicate bleached FRAP area. Scale bars are 5 μm, time is indicated (seconds).

**Video S4. Control FRAP analyses. Related to Figure 3**. Live time-lapse imaging of FRAP analyses of free GFP turnover in in whole ovaries (top), and of H2A-GFP turnover (bottom) in whole ovaries, in the early forming Bb in the nuclear cleft (left) and in the mature Bb (right). Red boxes indicate bleached FRAP area. Scale bars are 5 μm, time is indicated (seconds).

**Video S5. 3D SIM of the mature Bb. Related to Figure 3E**. 3D SIM super-resolution views of the mature Bb (Buc-GFP, green).

**Video S6. 3D microtubule organization in respect to the cleft condensate and the mature Bb. Related to Figure 4**. 3D oocyte views showing microtubule (α-tubulin, magenta) and the early (left) or mature (right) Bb condensate (Buc-GFP, cyan).

**Video S7. FRAP analyses of Buc in nocodazole- and DMSO-treated ovaries. Related to Figure 4. A.** Live time-lapse imaging of FRAP analyses of Buc-GFP turnover in the early forming Bb in whole ovaries, treated with control DMSO (left) and nocodazole (right). White outline indicates oocyte contour, red boxes indicate bleached FRAP area. Scale bars are 5 μm, time is indicated (seconds).

**Video S8. FRAP analyses of Buc in colchicine- and DMSO-treated ovaries. Related to Figure 4. A.** Live time-lapse imaging of FRAP analyses of Buc-GFP turnover in the early forming Bb in whole ovaries, treated with control DMSO (left) and colchicine (right). White outline indicates oocyte contour, red boxes indicate bleached FRAP area. Scale bars are 5 μm, time is indicated (seconds).

**Video S9. FRAP analyses of Buc in taxol- and DMSO-treated ovaries. Related to Figure 4. A.** Live time-lapse imaging of FRAP analyses of Buc-GFP turnover in the early forming Bb in whole ovaries, treated with control DMSO (left) and taxol (right). White outline indicates oocyte contour, red boxes indicate bleached FRAP area. Scale bars are 5 μm, time is indicated (seconds).

**Video S10. FRAP analyses of Buc in ciliobrevin- and DMSO-treated ovaries. Related to Figure 5. A.** Live time-lapse imaging of FRAP analyses of Buc-GFP turnover in the early forming Bb in whole ovaries, treated with control DMSO (left) and ciliobrevin (right). White outline indicates oocyte contour, red boxes indicate bleached FRAP area. Scale bars are 5 μm, time is indicated (seconds).

**Video S11. FRAP analyses of Buc in taxol-, ciliobrevin+taxol-, and DMSO-treated ovaries. Related to Figure 5. A.** Live time-lapse imaging of FRAP analyses of Buc-GFP turnover in the early forming Bb in whole ovaries, treated with control DMSO (left) and ciliobrevin (center), and taxol+ciliobrevin (right). White outline indicates oocyte contour, red boxes indicate bleached FRAP area. Scale bars are 5 μm, time is indicated (seconds).

**Videos S12. 3D analyses of Bb disintegration. Related to Figure 6**. Representative IMARIS 3D images of entire oocytes co-labeled for Buc (green, white arrowheads) and γTub (red, yellow arrowheads), and DNA (DAPI, blue), in control DMSO (top) and nocodazole (bottom) treated ovaries. White lines indicate the distance between automatically identified surfaces of Buc entities and the centrosome, as shown on the composite images (left) and without other channels (right). Red arrowheads point to nuclei of follicle cells that surround the oocytes, and are shown from different positions through the Z axis.

**Videos S13. Live time-lapse imaging of the mature Bb in nocodazole- and DMSO-treated oocytes. Related to Figure 7**. Live imaging of Buc-GFP oocytes from the experiments in Figure 7A-C. 3D IMARIS live images of entire oocytes are shown on the left, and zoom-in live images of the Buc-GFP labeled Bb are shown on the right. The video shows oocyte and zoomed in Bb images throughout the course of the experiment from the XY projection and then from YZ and XZ projections in control DMSO treated oocytes, followed by the corresponding images and projections in nocodazole treated oocyte. DNA (blue) is labeled by Hoechst. Scale bars are 10 μm (left) and 3 μm (right).

## Supplementary Figures

**Figure S1.**
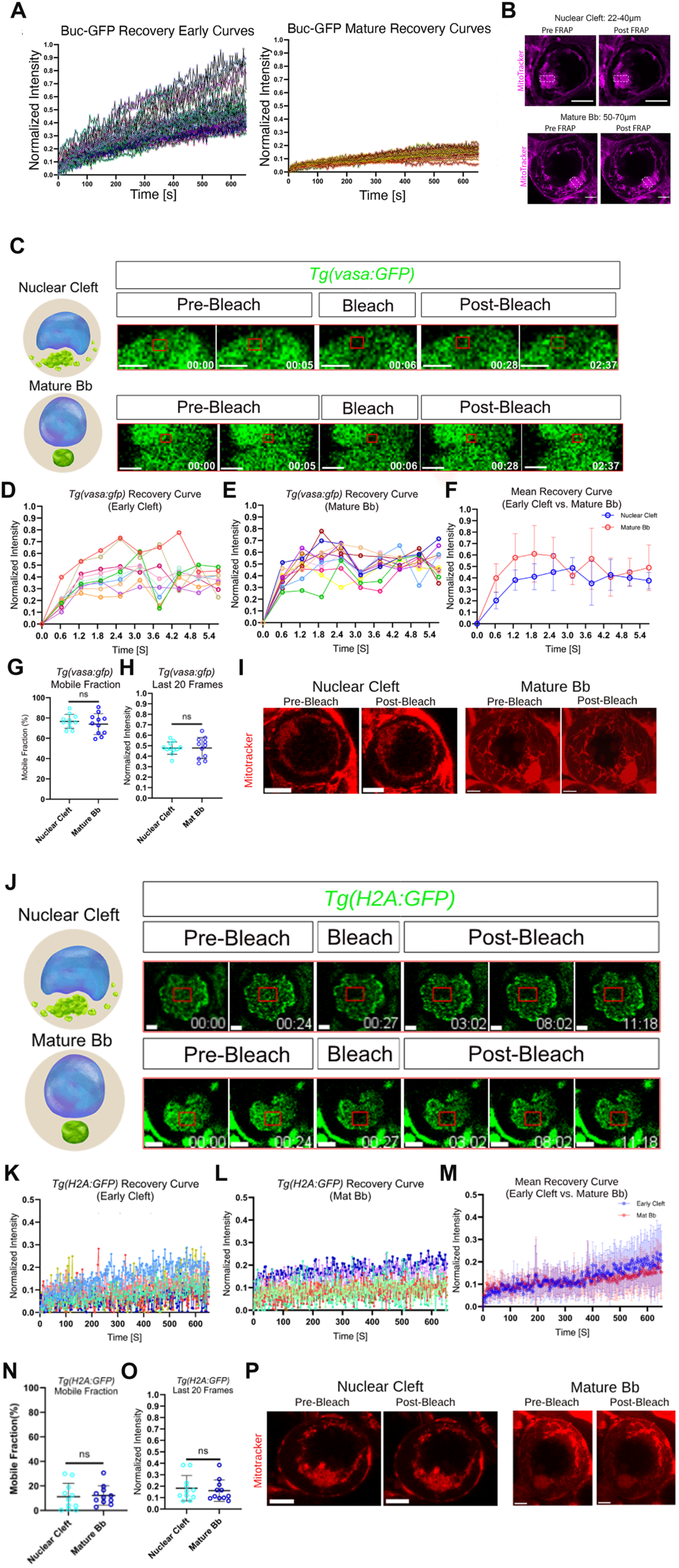
FRAP analyses for Buc -GFP and control proteins. Related to Figure 3. **A.** Individual Buc recovery curves of all oocytes (individual colors) in the FRAP experiments in Fig. 3A-E, at mature Bb (A) and early cleft (B) stages. **B.** Mitotracker labeling of live ovaries pre- and post-FRAP, in the FRAP experiments in Fig. 3A-E, shows viable active mitochondria after FRAP, as control for normal cell viability under our FRAP conditions. White dashed boxes indicate the bleached regions in the FRAP. Early cleft and mature Bb stages are shown. Scale bars are 10 μm. **C-I.** FRAP analysis of GFP in *vasa:GFP* ovaries, in the early forming Bb in the nuclear cleft (top) and in the mature Bb (bottom) stages. Images in C are snapshots at indicated timepoints (minutes) from representative Videos S4. Recovery plots for all individual oocytes early cleft (D) and mature Bb (E) stages, as well as pooled data for both stages (F) are shown. Quantification by mobile (G) fraction calculations, and by intensity at last 20 frames (H) are shown. Mitotracker labeling of live ovaries pre- and post-FRAP for both stages are shown in I. **J-P.** FRAP analysis of H2A-GFP in *h2a:H2A-GFP* ovaries, in the early forming Bb in the nuclear cleft (top) and in the mature Bb (bottom) stages. Images in J are snapshots at indicated timepoints (minutes) from representative Videos 4. Recovery plots for all individual oocytes early cleft (K) and mature Bb (L) stages, as well as pooled data for both stages (M) are shown. Quantification by mobile (N) fraction calculations, and by intensity at last 20 frames (O) are shown. Mitotracker labeling of live ovaries pre- and post-FRAP for both stages are shown in P. All bars are Mean ± SD. See also Videos S3-4.

**Figure S2.**
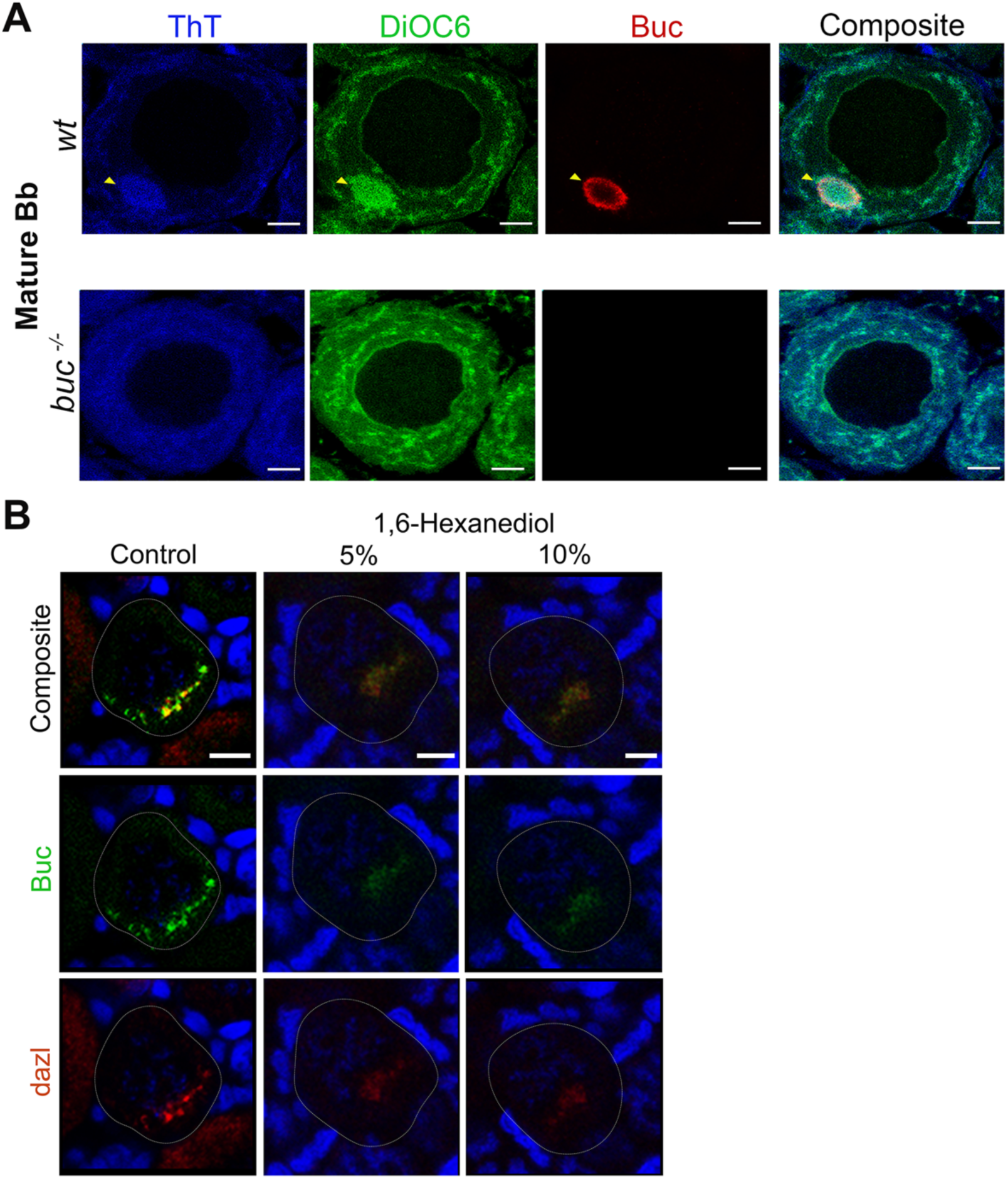
Presumptive amyloid β-sheets in the mature Bb and Hexanediol sensitivity of the early condensate. Related to Figure 3. **A.** Detection of presumptive amyloid β-sheets (ThT, blue, arrowheads) in the Bb in fixed wt ovaries (top) labeled for with Buc antibody (red), and the lipid dye DiOC6 (green), which detects Bb enriched mitochondria. In *buc^-/-^* ovaries (bottom) in which the Bb fails to form as evident by the loss of Buc and the loss of DiOC6 enrichment, ThT is not detected above background levels. Sale bars are 10 μm. **B.** Ovaries co-labeled for Buc (green) and *dazl* mRNA (HCR FISH, red) after 5% or 10% 1,6-Hexanediol treatment (middle and right panels, respectively), compared to control untreated ovaries (left). In all images, DNA (blue) is labeled by Hoechst (live) or DAPI (fixed) ovaries. Bright nuclei belong to somatic follicle cells, that surround oocytes. Scale bars are 10 μm.

**Figure S3.**
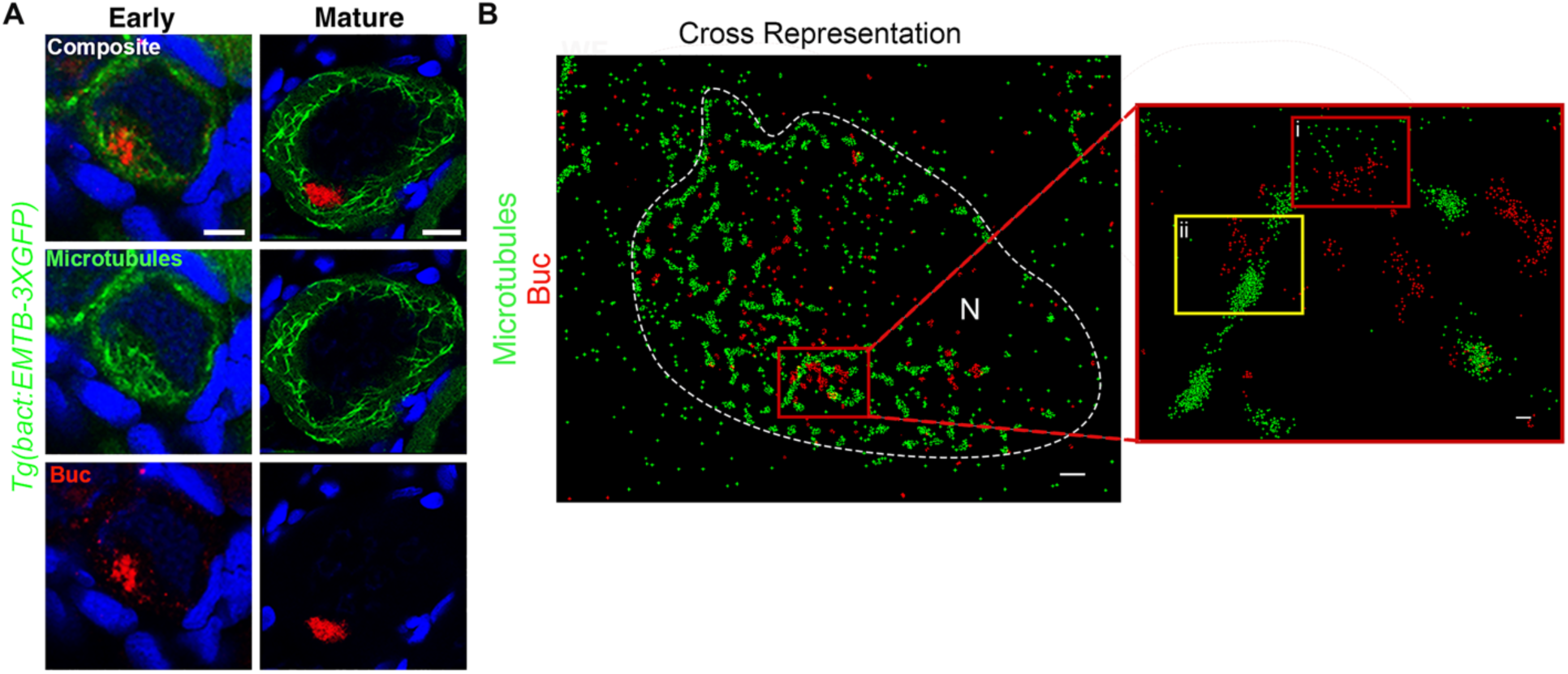
Microtubule organization with respect to Buc. Related to Figure 4. **A.** Co-dynamics of Buc (red) and microtubules (EMTB-GFP, green) during oocyte development at early and mature Bb stages. DNA (blue) is labeled by DAPI. Scale bars are 10 μm. **B.** Cross-visualization of the dSTORM super-resolution Gaussian images shown in Fig. 4C. Scale bars are 100 nm. N indicates the oocyte nucleus.

**Figure S4.**
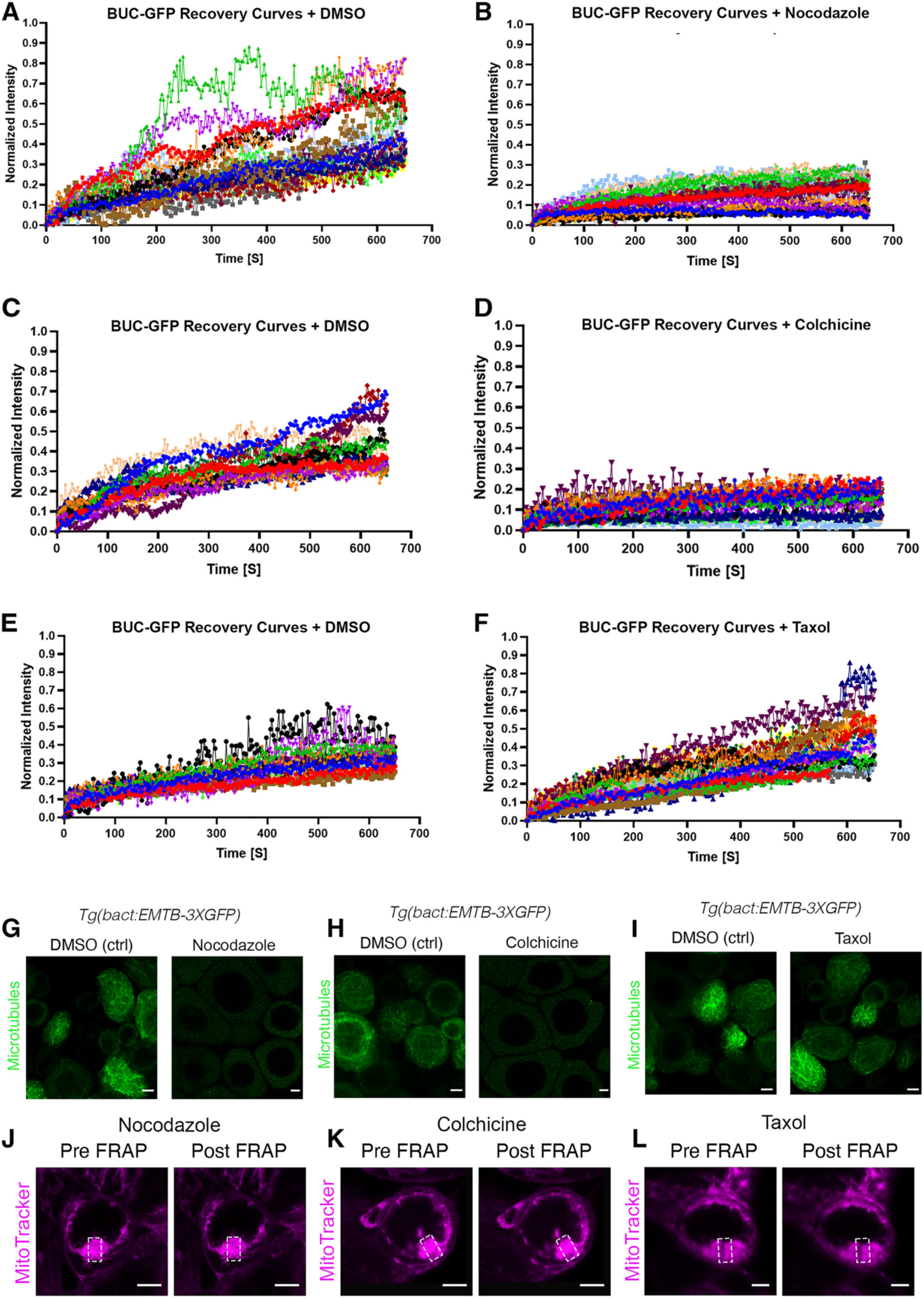
Raw data and controls for FRAP analyses. Related to Figure 4. **A-F.** Individual Buc recovery curves of all oocytes (individual colors) in the FRAP experiments in Fig. 4, in control DMSO (A, C, E) and drug treated (B, D, F) ovaries. **G-I.** Control EMTB-GFP ovaries treated side by side, show microtubule depolymerization upon nocodazole (G), and colchicine (H), but not taxol (I) treatments. **J-L.** Mitochondria and cell viability controls for FRAP experiments. White dashed boxes indicate the bleached regions in the FRAP. All scale bars are 10 μm. See also Videos S7-9.

**Figure S5.**
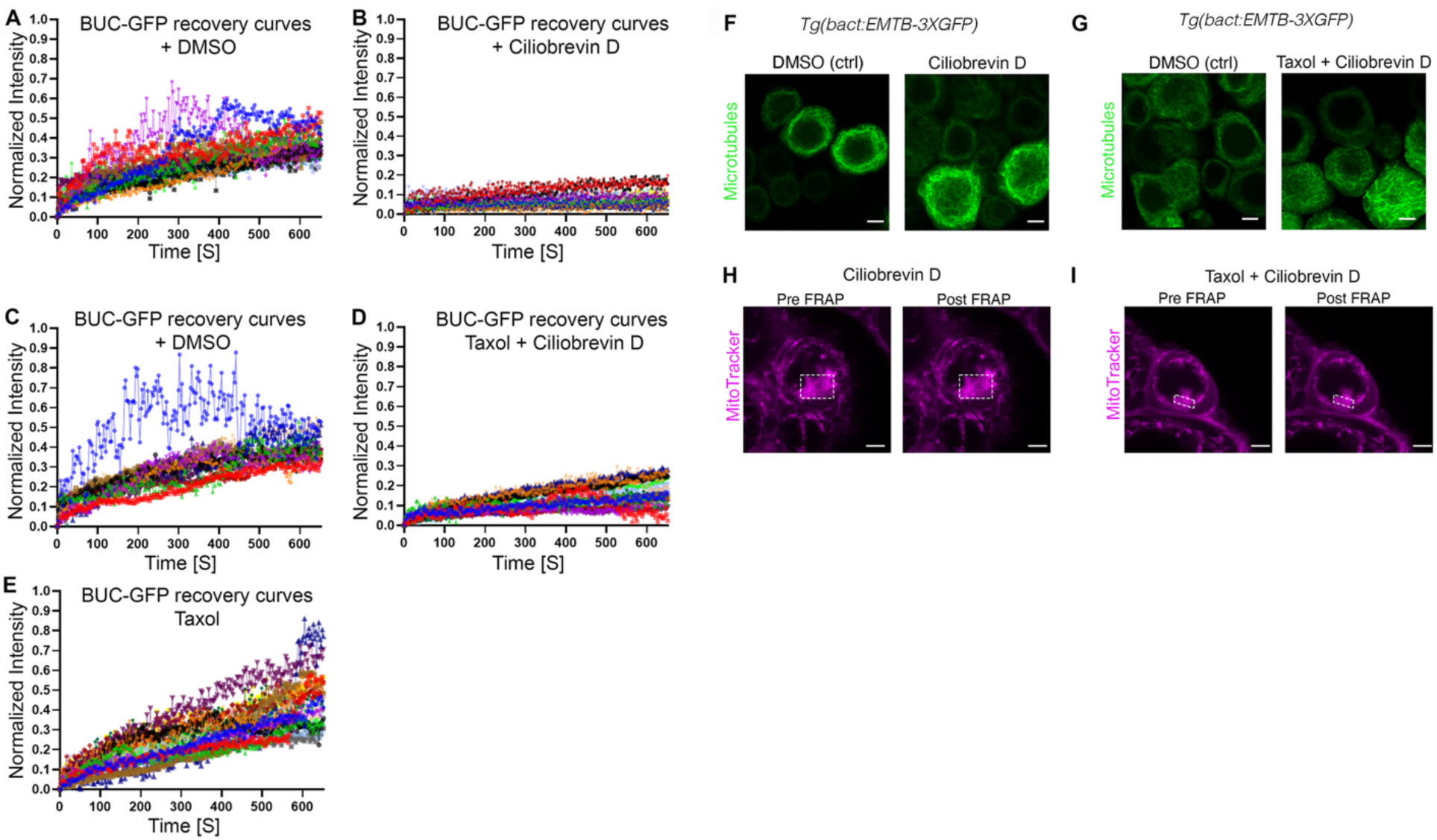
Raw data and controls for FRAP analyses. Related to Figure 5A-H. **A-E.** Individual Buc recovery curves of all oocytes (individual colors) in the FRAP experiments, in control DMSO (A, C) and drug treated (B, D, E) ovaries. **F-G.** Control EMTB-GFP ovaries treated side by side, show grossly intact microtubules with no apparent depolymerization upon ciliobrevin (F), and taxol+ciliobrevin (G) treatments. **H-I.** Mitochondria and cell viability controls for FRAP experiments. White dashed boxes indicate the bleached regions in the FRAP. All scale bars are 10 μm. See also Videos S10-11.

**Figure S6.**
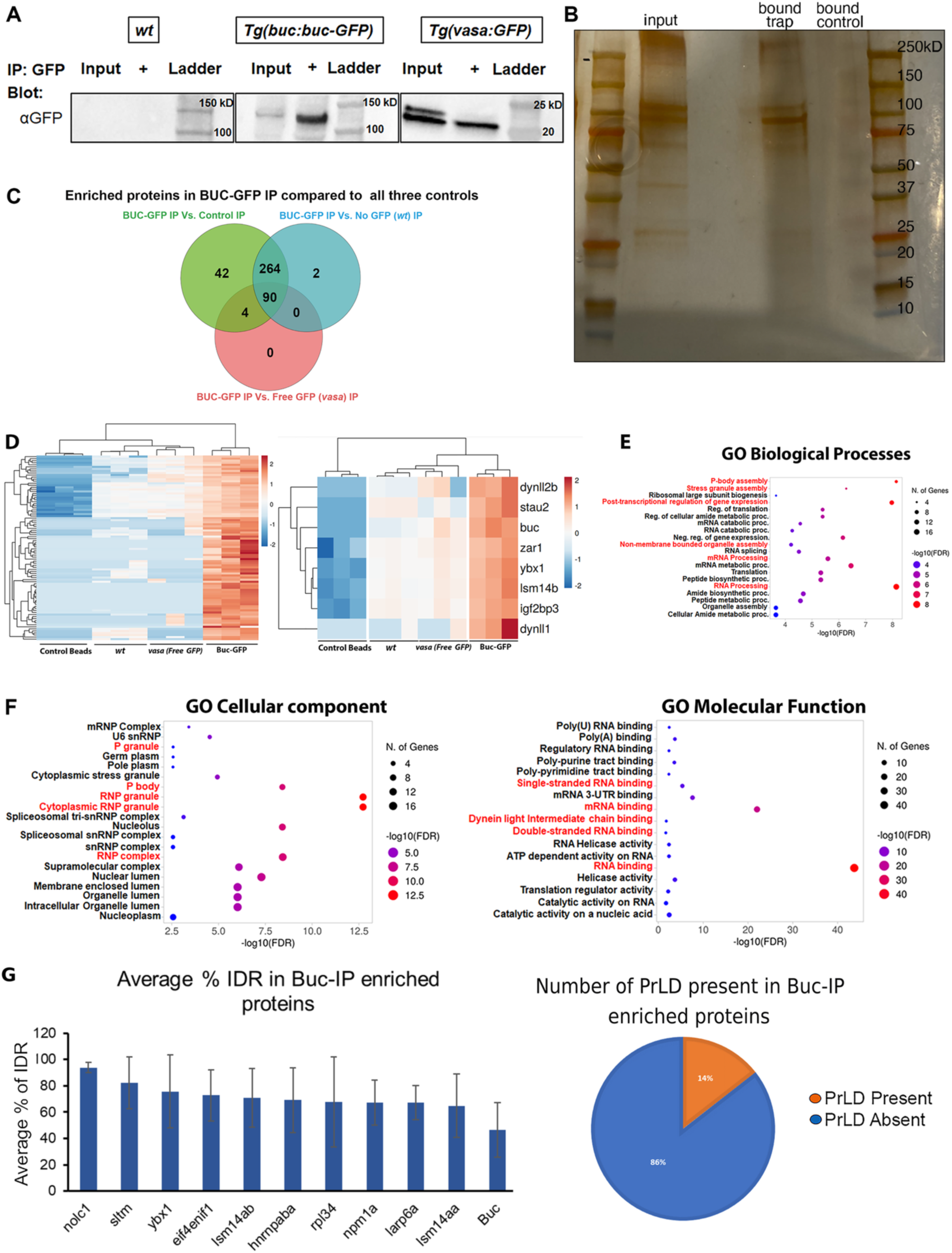
Controls, raw data and analyses of Buc-GFP IP-MS experiments. Related to Figure 5K-M. **A.** Western blot control for Buc-GFP-IP experiments. GFP proteins are not detected in either input or Buc-GFP-IP on wt samples (no transgenic GFP; left), but are detected in both input and IP samples from Buc-GFP-IP on Buc-GFP expressing ovaries (middle) and free GFP expressing ovaries (vasa:GFP; right). **B.** Raw silver-stain analysis shows detection of proteins in input samples from Buc-GFP expressing ovaries, as well as pulled down proteins from Buc-GFP-IP samples, but not in control samples (beads with no GFP nanobody). Since the Silver-stain reaction develops rapidly gel images were taken by smartphone and some shadowing reflection of the gel might be seen. **C.** Venn diagram showing comparisons between Buc-GFP-IP samples and each independent control in IP-MS experiments, showing 90 proteins that are specifically enriched against all controls. **D.** Heatmap and hierarchical clustering of the 90 proteins that are specifically enriched in Buc-GFP-IP samples against all three controls (left), including known Bb proteins, and Dynll1 and Dynll2b (right). **E-F.** GO analyses for Biological processes (E), Cellular components (F, left) and Molecular function (F, right) identifies relevant categories, including RNA processing, RNP-P body and stress-granule assembly, and non-membrane bound organelles (highlighted in red). G. Left: Top ten Buc-IP’ed proteins that contain substantial (∼65-85%) IDR sequences. Right: Percentage of Buc-IP’ed proteins with or without a predicted Prion-like domain (PrLD). See also Table S2.

**Figure S7.**
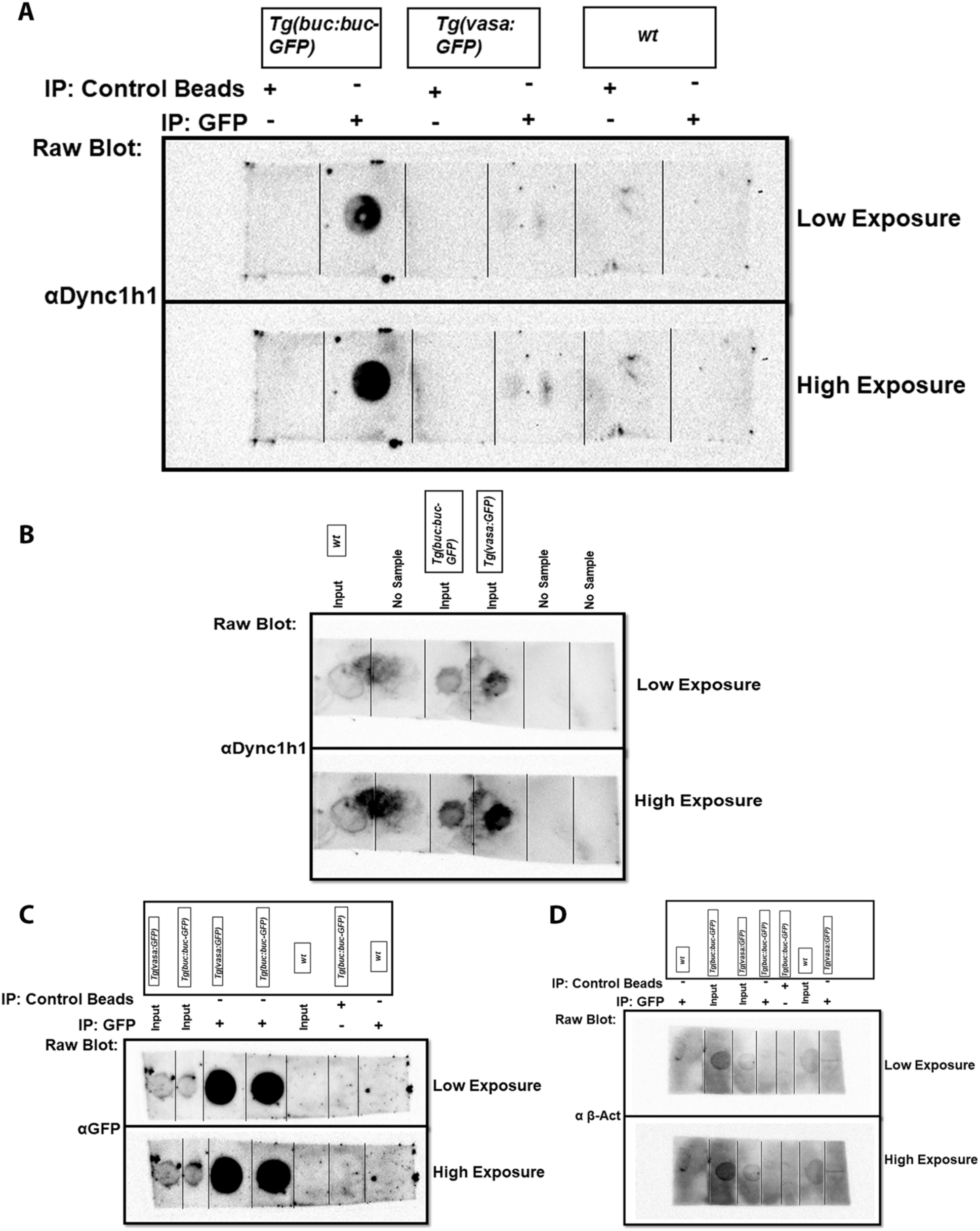
Raw blots of Buc-GFP IP of Dync1h1. Related to Figure 5N. Raw membranes for the dot-blot analyses of the Buc-GFP-IP experiment. **A.** IP samples blotted with Dync1h1 antibody. **B.** Input samples blotted with Dync1h1 antibody. **C.** IP and input samples blotted with GFP antibody. **D.** IP and input samples blotted with β-Act antibody.

**Figure S8.**
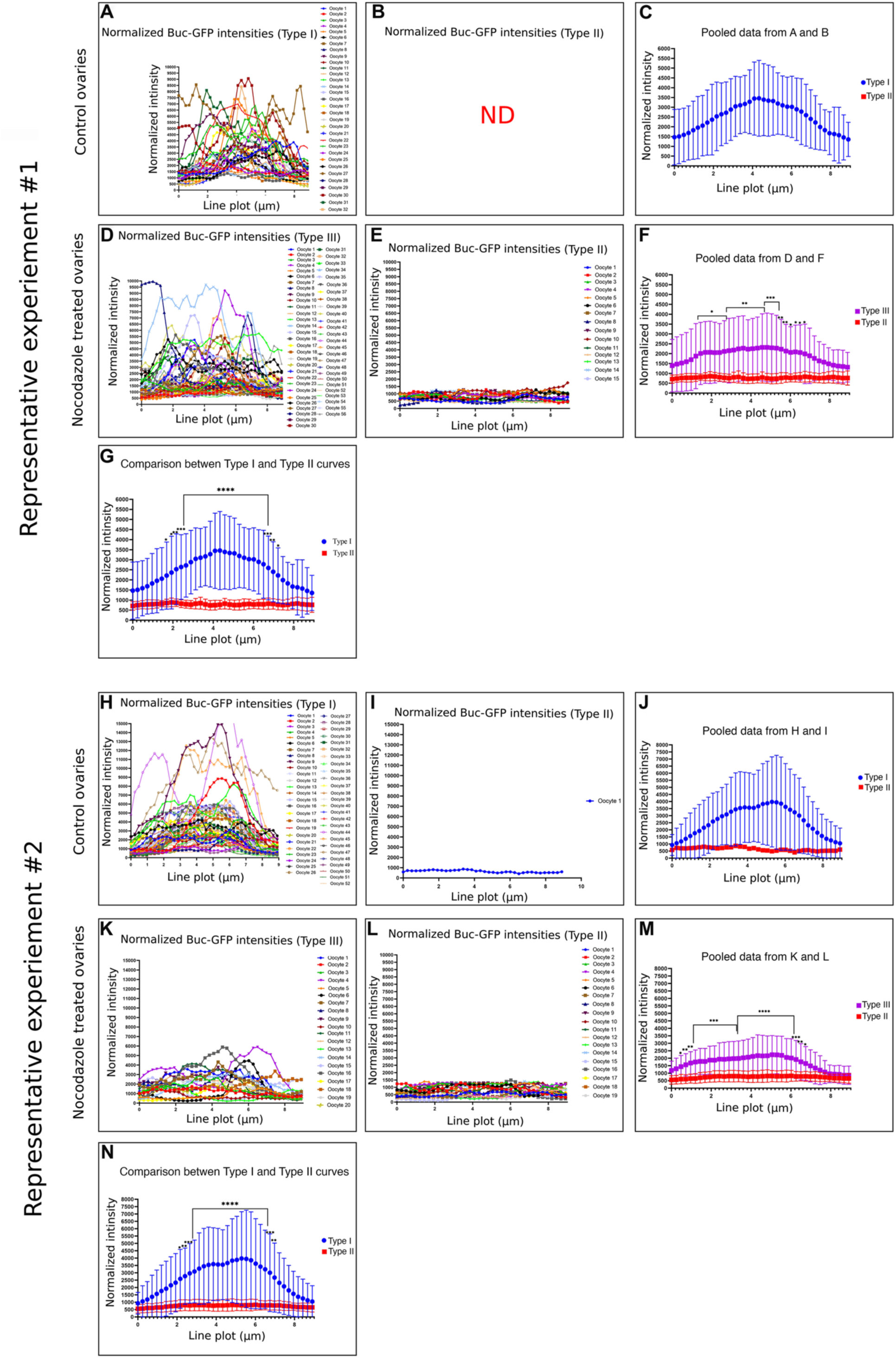
Raw data of Buc disintegration experiments. Related to Figure 6. Representative raw data for the experiments in Fig. 6D-J, using the centrosome (γTub) as a marker for the cleft vicinity, as shown in Fig. 6C. Two representative experiments are shown #1 and #2. For each experiment, raw normalized Buc intensity curves of all individual oocytes (individual colors) are shown in control DMSO (type I: A, H, type II: B, I), and nocodazole treated ovaries (type I: D, K, type II: E, L). Plots in C and J are pooled data (type I and type II) from DMSO control ovaries, and in F and M plots are similarly pooled data from nocodazole treated ovaries, in each experiment. Plots in G and N are comparisons between type I curves from control DMSO treated ovaries and type II curves from nocodazole treated ovaries, demonstrating they are statistically different. All bars are Mean ± SD.

**Figure S9.**
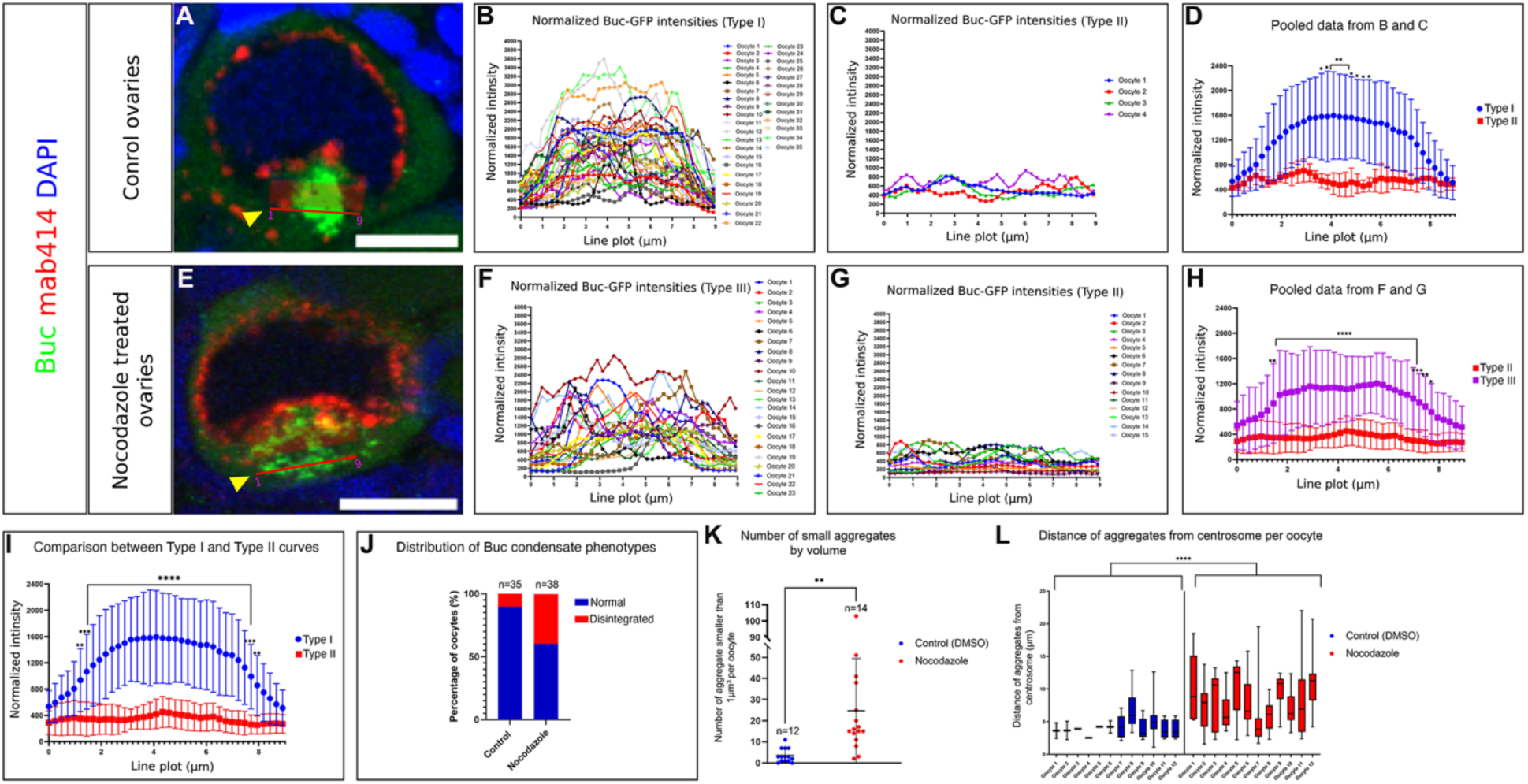
Raw data of Buc disintegration experiments. Related to Figure 6. Representative raw data for the experiments in Fig. 6D-I, using the mAb414 as a marker for delineating the cleft. **A, E.** representative images of DMSO control (A) and nocodazole treated (E) ovaries labeled with mAb4141 (red), Buc (green) and DNA (DAPI, blue). The Buc intensity measuring lines are indicated by arrowheads. Scale bars are 10 μm. **B, C, F, G.** Raw normalized Buc intensity curves of all individual oocytes (individual colors) are shown in control DMSO (type I: B, type II: C), and nocodazole treated ovaries (type I: F, type II: G). **D, H.** Plots in D are pooled data (type I and type II) from DMSO control ovaries, and in H plots are similarly pooled data from nocodazole treated ovaries, in each experiment. **I.** Plots are comparisons between type I curves from control DMSO treated ovaries and type II curves from nocodazole treated ovaries, demonstrating they are statistically different. **J.** Plots are the rate of each type I and type II curves in DMSO control and nocodazole treated ovaries in this experiment. All bars are Mean ± SD. **K.** the number per oocyte of granules with volume smaller than 1 μm^3^ for DMSO control and nocodazole treated ovaries are plotted. **L.** Raw data for Fig. 6O, plotting the distance between all granules to the centrosome in each oocyte, in the DMSO control and nocodazole treated ovaries. All bars are Mean ± SD. See also Video S12.

**Figure S10.**
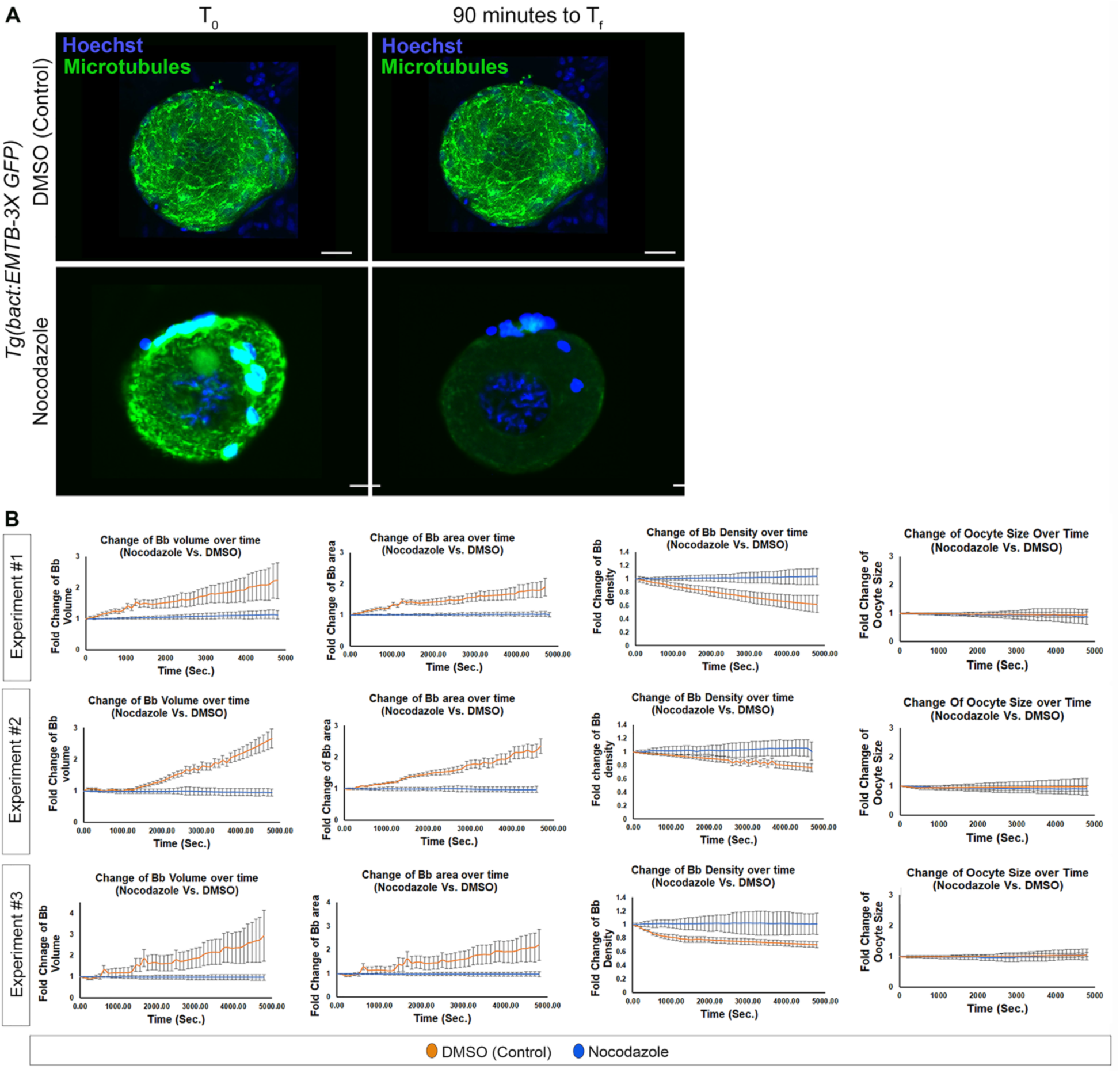
Controls and raw measurements from live imaging of the mature Bb upon DMSO or nocodazole treatments. Related to Figure 7. **A.** Representative live images of EMTB-GFP oocytes treated with DMSO control (top) and nocodazole (bottom), side by side to the experiments in Fig. 7A-C, at the beginning (T*_0_*) and end (T*_f_*) of incubation, showing intact microtubules in DMSO control and their depolymerization by nocodazole. DNA (Hoechst, blue) show oocyte nuclei as well as nuclei of surrounding follicle cells. Scale bars are 10 μm. **B.** Raw data for Fig. 7C from three independent experiments (top, middle, and bottom rows). For each experiment, the pooled data per experiment are plotted for fold-change over time of Bb volume (left), surface area (middle), and density (right), in DMSO control (blue) and nocodazole treated (orange) oocytes. All bars are Mean ± SD. See also Video S13.

